# Input-output signal processing plasticity of vagal motorneurons in response to cardiac ischemic injury

**DOI:** 10.1101/2020.08.09.242792

**Authors:** Jonathan Gorky, Alison Moss, Marina Balycheva, Rajanikanth Vadigepalli, James S. Schwaber

## Abstract

Vagal stimulation is emerging as the next frontier in bioelectronic medicine to modulate peripheral organ health and treat disease. The neuronal molecular phenotypes in the dorsal motor nucleus of the vagus (DMV) remain largely unexplored, limiting the potential for harnessing the DMV plasticity for therapeutic interventions. We developed a mesoscale single cell transcriptomics data from hundreds of DMV neurons under homeostasis and following physiological perturbations. Our results revealed that homeostatic DMV neuronal states can be organized into distinguishable input-output signal processing units. Remote ischemic preconditioning induced a distinctive shift in the neuronal states towards diminishing the role of inhibitory inputs, with concomitant changes in regulatory microRNAs miR-218a and miR-495. Chronic cardiac ischemic injury resulted in a dramatic shift in DMV neuronal states suggestive of enhanced neurosecretory function. We propose a DMV molecular network mechanism that integrates combinatorial neurotransmitter inputs from multiple brain regions and humoral signals to modulate cardiac health.

## Introduction

We aim to begin to unravel how the dorsal motor nucleus of the vagus (DMV) responds to physiological perturbations and interacts with the periphery via the vagus nerve. It is clear that the DMV is a central modulator of homeostatic function of multiple organ systems based upon the anatomy of projecting vagal neurons ^1–4^. The nucleus remains largely understudied as its integrative and functional capacity has been obscure, especially with regard to influence on the heart. However, recent findings suggest that DMV activity is critical to heart health and has the potential to rescue the heart from damage through its coordination of direct cardiac projections and indirect projections to the gut, which can be induced through remote ischemic preconditioning (RIPC) ^5^ ^6, 7^. Traditional understanding of neuronal function suggests mediation of effector functions primarily through selective neuronal activity of one neuron type or another. Therefore, most attempts at characterizing the heterogeneity within the transcriptional landscape of neurons made the basic assumption that these neuronal subtypes were essentially fixed within the timescale of physiological perturbations and thus a snapshot in time was sufficient to characterize them ^8^. Subsequent work has since shown shifts in these landscapes in response to development, aging, or even caloric restriction, showing that dynamic changes from one neuronal state to another were not only possible, but were a matter of course in normal physiology ^9, 10^. We formulate our approach in light of these findings to examine the potential shift in DMV neuronal state in response to acute and subacute physiological perturbations including cardiac ischemic injury.

The shifting of DMV neuronal states in response to physiological perturbations likely serves to alter a neuron’s effector function (an ‘output’) and/or its ability to be influenced by a projection (i.e., an ‘input’) ^11^. Hence, a useful way to delineate neuronal states is by considering each state as representative of a particular type of signal processing unit based on combinatorial weighting of a class of inputs (receptor expression) and unique collection of a class of outputs (neurotransmitters and peptides). To investigate the distribution of such signal processing units in the DMV, we performed high-throughput microfluidic RT-qPCR of laser captured single neurons and small pools of less than 5 neurons, to develop a targeted mesoscale gene expression data set with high sensitivity, specificity, and replicability ^12, 13^. In each single cell scale sample, we profiled a large panel of neuronally relevant genes including signaling pathways, high-yield receptors, neurotransmitter enzymes, and neuropeptides collated from a wide-ranging survey of the literature on DMV gene and protein expression, and neuronal connectivity (Supplemental File 1). We sought to analyze the data for gene expression modules and gene coexpression correlation networks, and organized the results into distinct input-output signal processing units that represent the potential interaction strength of inputs from several brain regions to the DMV and the combinatorial effector molecules as putative outputs of these neuronal states (Fig. 1a). We examined the potential shift in these DMV neuronal states in response to acute physiological perturbation of remote ischemic preconditioning (RIPC), whose effects on the heart occur within hours and require alteration of DMV neuronal activity ^5–7, 14^. In addition, we induced longer timescale dynamic changes in the DMV neuronal landscape through ligation of the left anterior descending (LAD) coronary artery in a chronic myocardial ischemia model over the course of 3 weeks. The observed changes in the distribution of neuronal states are representative of alterations in the input/output signal processing function of the nucleus as a whole and specific to each physiological perturbation. Our proposed framework to analyze DMV neuronal state landscape as a set of input-output signal processing units affecting neuromodulatory action of vagal outflow offers new insight into how the DMV accomplishes its autonomic regulatory function by dynamically responding to the demands of the organismal physiology.

**Fig. 1:**
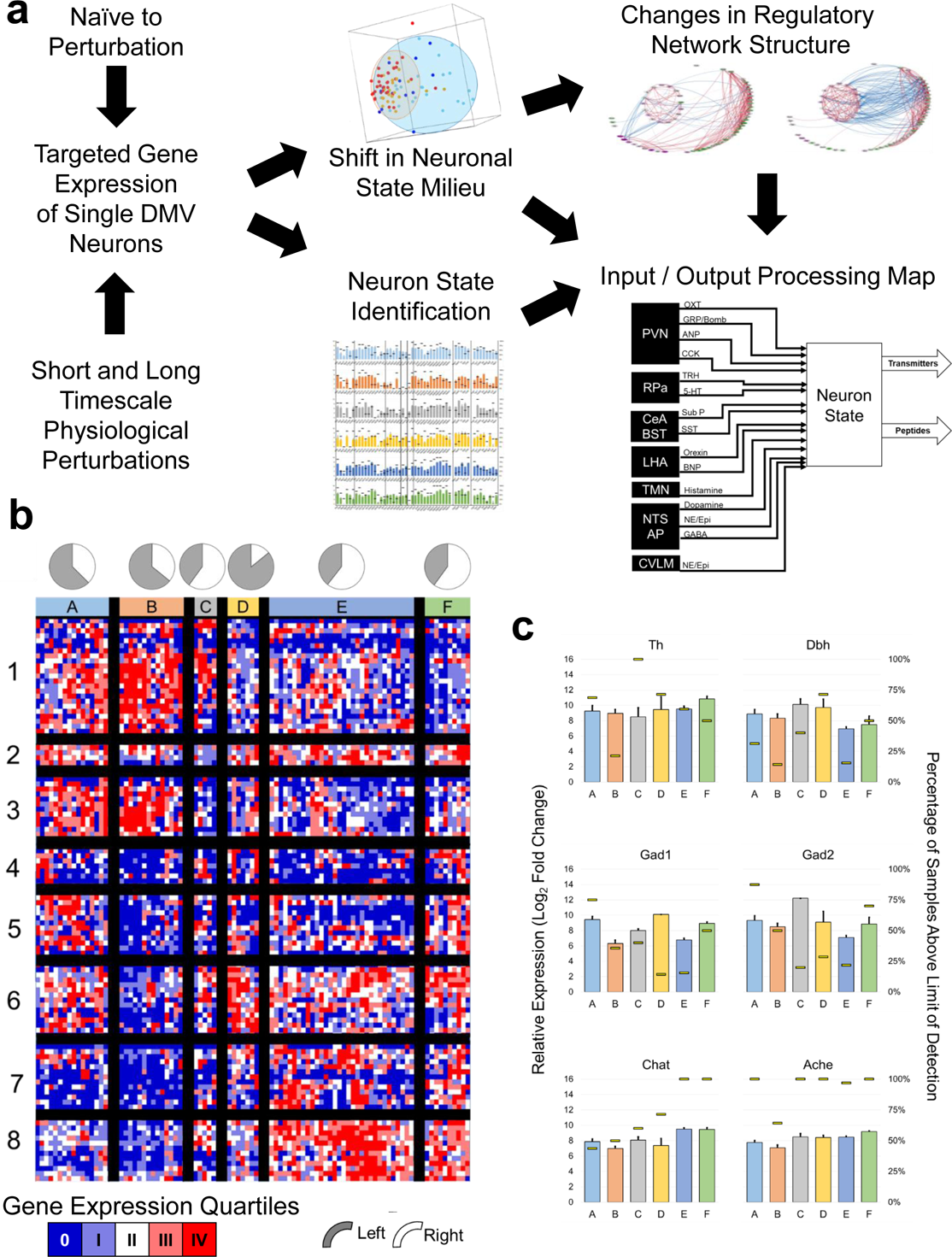
Single cell gene expression profiling delineates distinct neuronal states in the dorsal motor nucleus of the vagus (DMV). **a** Experimental and analytical workflow to determine the DMV neuronal states and their shifting responses to physiological perturbations. The single neuron scale gene expression data was analyzed to identify correlational neuronal clusters corresponding to distinguishable molecular states. These states, taken together with the correlated networks of receptors and neurotransmitters/neuropeptides, were used to infer state-specific input-output signal processing maps connecting DMV neuronal phenotypes to upstream brain regions as well as delineating downstream neuromodulatory systems. **b** Heatmap showing six distinct neuronal states (A-F) based on hierarchical clustering of single cell scale expression of genes corresponding to neuropeptide/neurotransmitter production and neuropeptide receptors. Data colored to represent the quartiles of gene expression values with dark blue showing values below the limit of detection. Pie charts show the proportion of samples within each state corresponding to left versus right DMV. **c** Neuronal state-wise expression of typical neurotransmitter systems conventionally used to delineate central neuronal phenotypes. Colored bars indicate expression level relative to the limit of detection (mean +/- SEM, left axis). Yellow horizontal lines show the percentage of samples with expression above the limit of detection within each state (right axis). The state-wise expression patterns of additional genes of interest are shown in the supplemental figures: neuropeptides (Fig. S2), neuropeptide receptors (Fig. S3), and calcium channel subunits (Fig. S4).

## Results

### Characterization of homeostatic molecular states of DMV neurons

We started with delineation of DMV neuronal states in the homeostatic state devoid of any specific physiological perturbations. We obtained a single neuron scale data set from 178 neurons and assayed 169 genes in each sample. Hierarchical clustering of quartile binned data yielded six transcriptional phenotypes (Fig. 1b; Supplemental Table 1 contains the details of the associated gene modules). These six DMV neuronal states were not distributed along the lines of canonical neurotransmitter production, as several neurotransmitter systems were abundantly expressed in multiple neuronal states (Fig. 1C). However, peptides (Fig. S1) ion channels (calcium channels highlighted in Fig. S2), and peptide receptor subtypes (Fig. S3) showed differential expression across the six DMV neuronal states, suggesting differential neuromodulatory functions.

We analyzed the six homeostatic neuronal states to develop a signature set of genes that are informative of the neuromodulatory signal processing within each state. We assessed a wide range of literature to construct an input-output signal processing structure onto which we mapped the differential gene expression data to infer distinctive neuromodulatory functions of DMV neurons (Supplemental Text). In this scheme, the relative differential expression pattern of genes corresponding to the processing of afferent signals (e.g., receptors) and those of effector functions (e.g., neurotransmitters) indicates the input-output signal processing likely to occur in the DMV neurons in a given state (Fig. 2a; Fig. S4). We represented the gene expression signatures corresponding to the six neuronal states (Fig. 1b) as distinguishable input-output signal processing units (Fig. 2b). Neuronal states A and F mainly correspond to the GABAergic phenotypes (Fig. 2b; Figs. S5,S6,S7, and S8) with state A including mostly *Gad*+ subtype G_A with a little bit of G_B and F including subtypes G_C and G_D. In neuronal states B and E there is a small cohort of *Gad*+ neurons that share some characteristics of G_B, but not all (e.g. expression of *Chat*). Further examination of Gad+ neurons in the DMV can be found in the supplemental text.

**Fig. 2:**
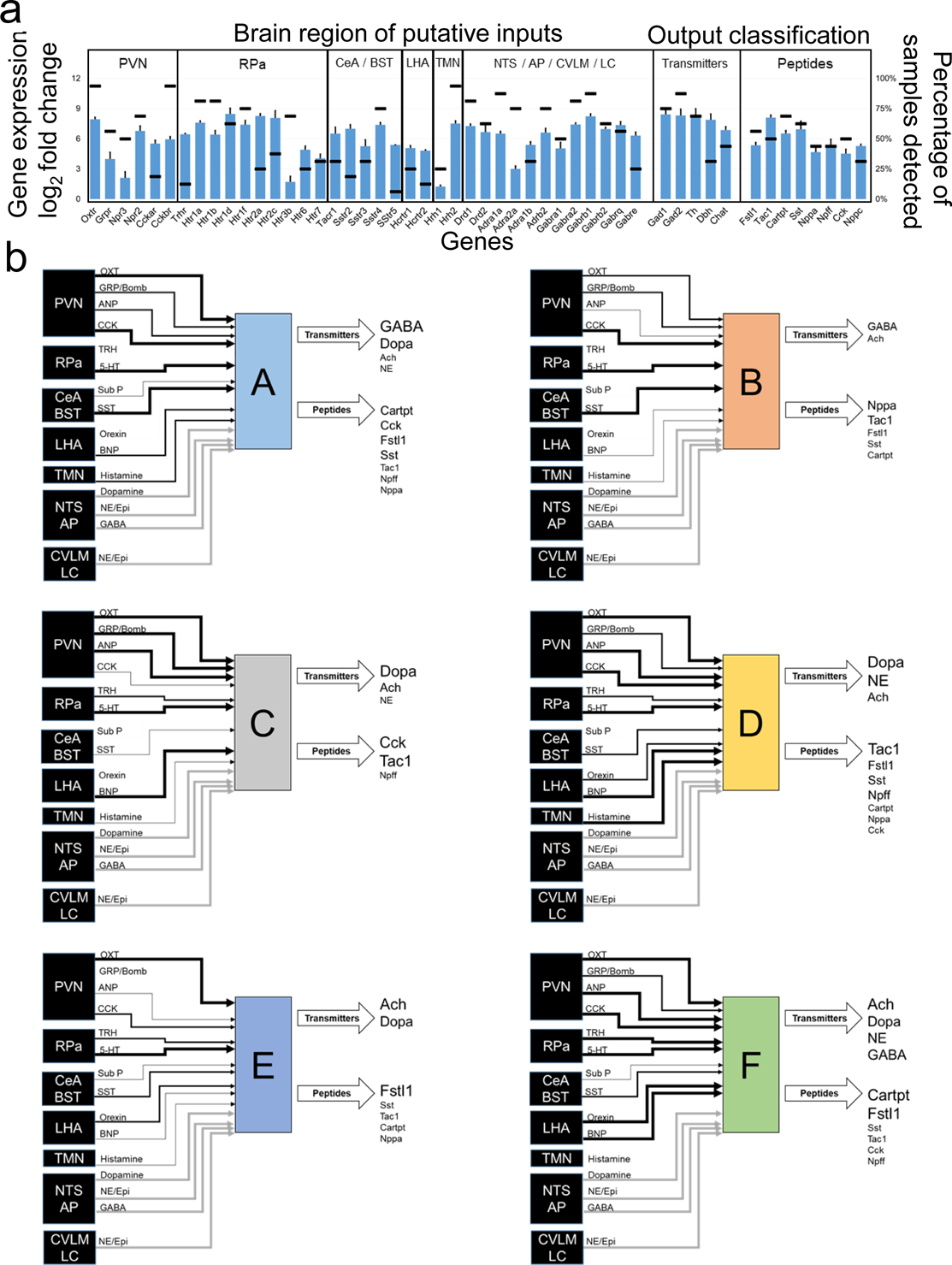
Input output signal processing map of DMV neuronal states. **a** Gene expression profile of DMV neurons in state A indicating the levels of a range of receptors linking these neurons to other brain regions (input signals) and the neurotransmitter/neuropeptide systems expressed in these neurons (output signals). Several brain regions projecting onto DMV neurons utilize specific neuropeptidergic systems, enabling inference of input connectivity and putative interaction strength based on receptor expression level within DMV. Detailed methods for network inference and supporting literature are included in the methods. Bar graph corresponds to the expression level relative to the limit of detection (left axis) and the horizontal lines correspond to the percentage of samples above the limit of detection for each gene (right axis). The input-output gene expression profiles of other states B-F are shown in the Supplementary Fig. S1. **b** Network representations of the input-output signal processing map of DMV neuronal states. The thickness of lines connecting the brain regions to a specific DMV neuronal state is proportional to the gene expression level of the receptors corresponding to the neuropeptide inputs, signifying putative interaction strength and information flow from these brain regions into DMV. The neurotransmitter/neuropeptide output signals from the DMV neurons are indicated in varying text size proportional to the gene expression level of corresponding enzymes. Within each DMV neuronal state-specific map, the input and output signals corresponding to the genes with expression below the limit of detection in those neurons are not shown. The regions NTS, AP, CVLM, and LC are grouped together as the corresponding inputs through dopamine, GABA, and norepinephrine cannot be specifically assigned to one of the regions. These inputs are shown in gray for all states. PVN - Paraventricular Nucleus; Rpa - Raphe Pallidus; CeA - Central Amygdala; BST - Bed Nucleus of the Stria Terminalis; LHA - Lateral Hypothalamus; NTS - Nucleus Tractus Solitarius; TMN - Tuberomammilary Nucleus; AP - Area Postrema; CVLM - Caudal Ventrolateral Medulla; LC - Locus Coeruleus; Ach - Acetylcholine; Dopa - Dopamine; NE - Norepinephrine (Noradrenaline); Epi - Epinephrine (Adrenaline); LC - Locus Coeruleus.

State B neurons have the highest expression levels of peptide receptors as well as most of the serotonin, dopamine, and histamine receptors. There were two distinct sub-states with a notable difference in expression of *Gad1* and *Gad2*. It is possible that given the diversity of more highly expressed receptors, these neurons play an integrative role taking a wide diversity of input. The production of several peptides in the relative absence of the neurotransmitter enzymes assayed here are suggestive of a peptidergic neurosecretory phenotype of neurons in state B.

Neuronal state E represents what may be considered a canonical efferent motor neuron from the DMV. These neurons express the milieu of channels that were observed by Goldberg and Gourine ^15, 16^ and have the requisite *Chat* and *Ache* expression corresponding to cholinergic neurons. These neurons also highly express the more standard GABAA and GABAB receptor subunits, in line with observations that a subset of DMV motor neurons were especially sensitive to GABAergic signaling ^17^. It is notable that neuronal state E constitutes a greater proportion of samples than any other of the distinguishable states. The relative lack of other receptors and the ion channels related to firing canonical action potentials rather than tonic or burst pacemaking (HCN and low voltage calcium channels) suggests that the unipolar neurons described in the monosynaptic gastric reflex circuits by Rinaman et al. may be primarily of this phenotype^18^.

Neuronal states C and D are most notable for their dominant expression of either gene group 1 or gene group 6 (Supplemental Table 1). Both clusters are made up of a relatively small number of samples, but have distinct gene expression patterns compared with nearest neighbor states B and E. State C neurons expressed much of the same canonical satiety peptides and neurotransmitter receptors as that of state B neurons, but without inputs derived from the same peptidergic neurons. The receptor expression profile in state B neurons suggests a greater interaction strength to respond to opioids, somatostatin, and cytokines like leukotriene B4 and IL-6 inputs. Neuronal state D is similar to state E, but with a more distinct upregulation of genes in group 6, whose membership includes multiple genes associated with pacemaker-like behavior (Hcn2, Girk2).Also of note is distinct upregulation of Hrh3 coding for the histamine 3 receptor, which is a moulatory hub of satiety in the rat ^19^.

### DMV neurons show specific state shifts in response to surgery and remote ischemic preconditioning

We obtained a comparable single neuron gene expression profiling data set on DMV neurons after RIPC (Fig. 3a). We compared the shift in DMV neuronal state due to RIPC to that in the homeostatic condition as well as due to sham surgery. Surprisingly, sham surgery had a prominent effect on gene expression in DMV neurons, even after a mere two and half hours. Several clusters differed from baseline in their sample distribution, most prominently the states A2, C and D (Fig. 3a). Both A2 and C have an over-representation of sham samples whereas D has an over-representation of RIPC samples. The main differences from A1 to A2 can be found in parts of gene modules 1,2, and 7 (Supplemental Table 1). The differences between the otherwise similar neuronal states C and D can be found mostly in gene cluster 1 which contains genes involved in sensing peptide and canonical neurotransmitters as well as key intracardiac neuron effectors like ANP and tachykinins as discussed later. These results indicate that neuronal response to RIPC leads to a DMV-wide gene expression pattern more similar to homeostatic naive samples rather than sham surgery, suggesting a RIPC-induced repression of the stress response seen in sham.

**Fig. 3:**
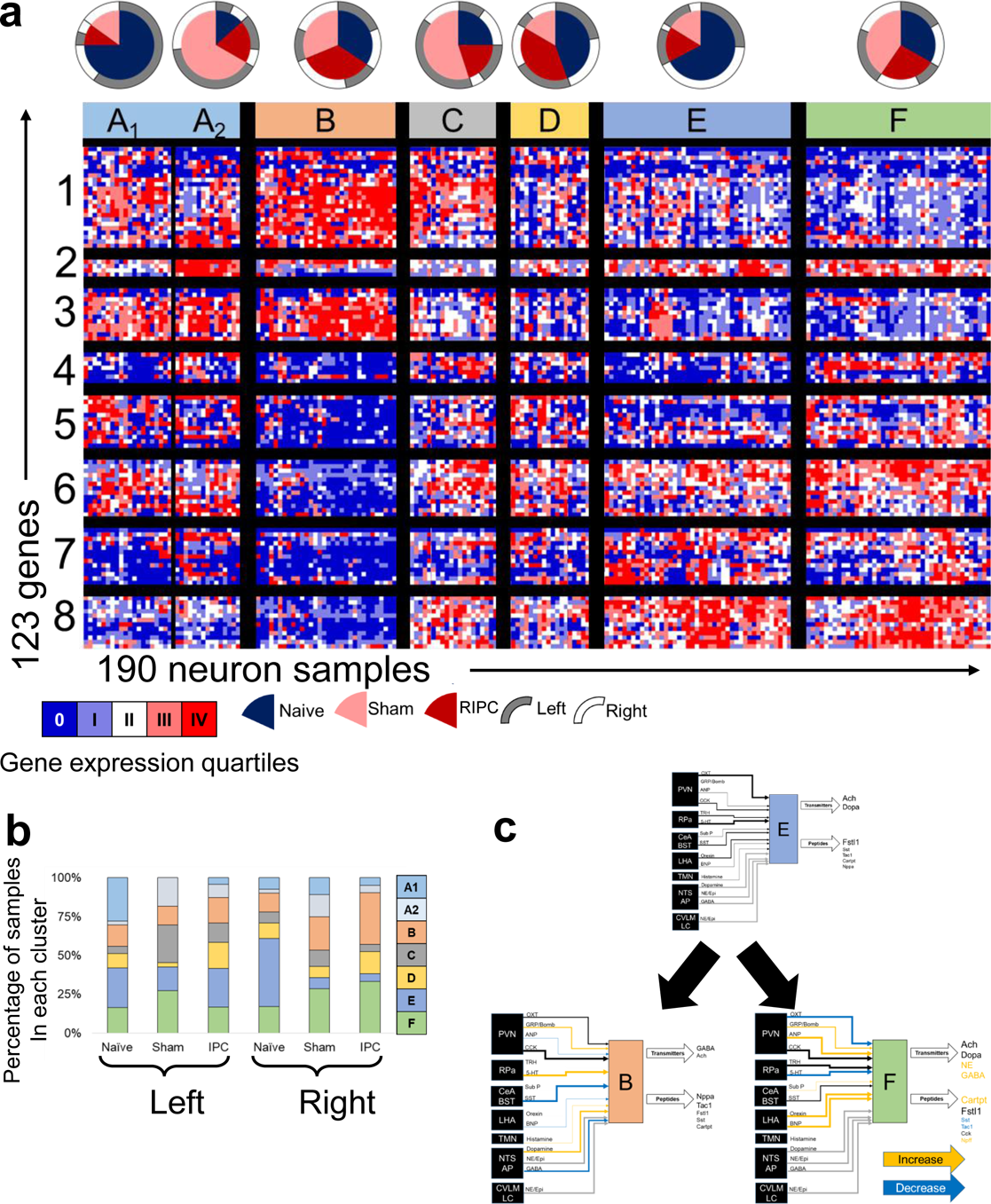
Physiological perturbations such as remote ischemic preconditioning (RIPC) and sham surgery shift the distribution of DMV neuronal states 2.5 hours. **a** Heatmap of single neuron gene expression showing the distribution of DMV states in homeostatic (naïve), sham surgical, and RIPC conditions. The proportion of neurons from each experimental condition is indicated in the pie charts. The border of the pie charts corresponds to the proportion of cells from left versus right DMV. Expression of a select subset of genes in a subset of states are illustrated in Fig.s 4a and 4b. **b** Distribution of neuronal states on the left and right sides of the DMV in the naïve, sham, and RIPC conditions. **c** Distinct neuronal states are enhanced in the right DMV after RIPC versus naive, evidenced by diminished state E with enhancement of state B and F. The physiological perturbation-induced changes in the gene expression of receptors (inputs) and neurotransmitter/neuropeptide systems (outputs) are mapped onto the state-specific networks as putative alteration of the interaction strength of input information flow or output signals: orange (increasing), black (unchanged), and blue (decreasing). PVN - Paraventricular Nucleus; Rpa - Raphe Pallidus; CeA - Central Amygdala; BST - Bed Nucleus of the Stria Terminalis; LHA - Lateral Hypothalamus; NTS - Nucleus Tractus Solitarius; TMN - Tuberomammilary Nucleus; AP - Area Postrema; CVLM - Caudal Ventrolateral Medulla; LC - Locus Coeruleus; Ach - Acetylcholine; Dopa - Dopamine; NE - Norepinephrine (Noradrenaline); Epi - Epinephrine (Adrenaline); LC - Locus Coeruleus.

In addition, the effect of the surgical treatment (RIPC or sham versus naive control) was significant in the *Fos*+ neuronal subset (Supplemental Text). The most prominent shift was the significant upregulation of *Th* in DMV neurons from the RIPC and sham cohorts (ANOVA, p<0.001), suggesting that DMV neurons can act like sympathetic neurons in an acute stress response.

A subset of DMV neurons appear to respond to RIPC or sham surgery by shifting to a novel state characterized by altered input-output signal processing structure reflected in the gene expression profile. This occurred in the sham condition most prominently on the left side of the DMV, where state A_1_ (the same as state A in Fig. 1b) shifted to the novel state A_2_ (Fig. 3b). The state transition of DMV neurons within the six signal processing structures delineated in Fig. 2b was readily apparent in the RIPC condition, particularly on the right side of the DMV where the decrease in proportion of neuronal state E coincided with an increase in the proportion of states B and F (Fig. 3b). We inferred the functional implications of such a shift in neuronal states by mapping the gene expression profiles to the input signal processing that is likely to be dialed up or down and which output effector functions are enhanced or suppressed. Our results suggest that following RIPC, the processing of inhibitory signals from the NTS and CeA are likely diminished, while at the same time amplifying the processing of excitatory signals from the RPa leading to an increased effect of natriuretic peptide A and tachykinins on the DMV through enhanced neuronal state B (Fig. 3c). By contrast, the sham surgery induced shift towards neuronal state F represents a potential amplification of response to excitatory inputs from LHA and PVN via a canonical sympathetic combination of neurotransmitter signals (NE and Cartpt) (Fig. 3c).

The RIPC-specific effects on DMV neuronal gene expression were subtle with the most notable change being a shift in histamine receptor subtypes (Fig. S10), suggesting increased interaction strength from the TMN. Gabra and Adra1a have a uniquely lower expression after RIPC, suggesting that RIPC diminishes inhibitory input signal processing mediated by GABA or NE (Fig. S10b). In addition, there is a reduction in expression of *Npr2* (signal transducer of natriuretic peptide) and up-regulation of *Npr3* (clearance receptor for the natriuretic peptide) in the DMV neurons (Fig. S10b). This shift in balance suggests a reduced influence of natriuretic peptide inputs from PVN to DMV, or alternatively, decreased influence from inhibitory interneurons producing natriuretic peptide within DMV. Also notable in the RIPC-specific response is a decrease in the number of DMV neurons expressing *Fstl1*. The cardioprotective peptide follistatin-1 can potentially be released via projections from DMV onto cardiac ganglia^20^ ^21^ and is known to suppress afferent transmission by potentiating Na+/K+ ATPase in neurons and myocytes ^22^. Our results suggest that the cardioprotection by RIPC reported by physiological studies ^23^ ^7^ ^5^ may not involve follistatin-1 mediated processes stimulated by vagal efferents originating from DMV.

### DMV microRNA regulatory network changes in response to cardiac ischemic injury and RIPC-mediated cardioprotection

With coordinate gene expression changes, it may be possible to determine likely effectors, such as microRNAs. Assaying microRNA changes in the DMV during LAD ligation with and without RIPC through find several that shift significantly in RIPC or that are renormalized by it (Fig. 4, S20, S21). Based on the p-value from template matching, fold change between LAD and AMC, and overall abundance, 3 microRNAs were identified as potential candidates for therapeutic intervention, miR-218a, miR-495, and miR-183. Of these 3 microRNAs, miR-495 has been shown to have strong cell type specificity, being heavily enriched in neurons compared to astrocytes, oligodendrocytes, and microglia ^24^. Of the top ten microRNAs that were determined to most likely affect the genes examined that were down regulated in RIPC (Fig. 4a), miR-495 was shown to actually have differential regulation during LAD ligation and be returned to normal levels with RIPC prior to LAD. Further examination of the gene network that these 3 microRNAs may be regulating was carried out using the miRWalk database. Genes with the highest binding prediction were taken into consideration. Genes that did not have robust expression in the DMV were filtered out using available RNAseq data in the brainstem. The remaining network represents the important genes involved in inflammatory and immune processes, excitatory and inhibitory receptors, ion channels, neuronal peptides and regulators, as well as transporters that are targeted by miRs-218a, 495, and 183 (Fig. 4d).

**Fig. 4:**
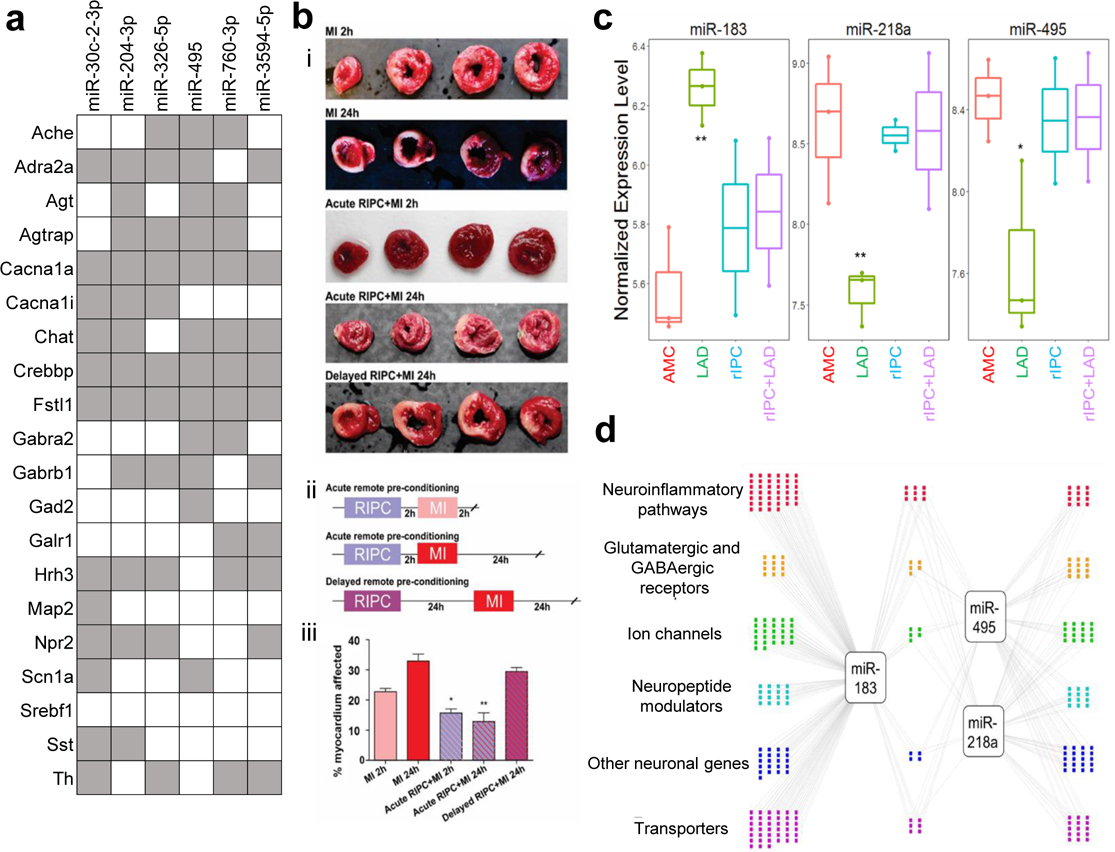
Likey microRNA mediators of DMV neuronal gene expression changes from RIPC. **a** Putative regulatory Interaction Matrix microRNA regulators RIPC-downregulated target genes While the significant genes that were discussed in this work are present, several other genes with the same expression pattern that did not meet the cut-off for statistical significance were also included in the query. Putative binding was determined through the use of several algorithms (mirWalk, RNA22, miRanda, and Targetscan) and a “hit” taken when more than one of these determined there to be a putative binding site in the 3’UTR. **b** Effects of remote ischemic preconditioning (rIPC) on the extent of myocardial tissue damage are present after 2 hours and subside by 24 hours. (i) Cardiac tissue after each treatment combination stained with triphenyl tetrazolium chloride (TTC) with areas of infarct shown in white and viable myocardium in red. (ii) Timeline of each RIPC experimental treatment group. The MI 2h and MI 24h groups did not have a rIPC component and only included induction of MI with the respective waiting time to tissue collection. (iii) Quantitative analysis of infarct size expressed as percentage of left ventricle. N=3-5 per group. *P<0.05, **P<0.005. **c** Putative microRNA control points in the DMV that showed dysregulation by LAD and normalization by rIPC. **P<0.005; *P<0.05, compared to age-matched control (AMC). We tested for expression and differential regulation of ∼400 microRNAs using nanoString digital counting platform. A total of 146 microRNAs were detected. Template analysis identified a subset of microRNAs that showed dysregulation in LAD and normalization by rIPC. Of these, a set of three microRNAs showed >2-fold differential regulation due to LAD compared to RIPC. **d** Differentially expressed microRNAs target genes in processes relevant to neuronal plasticity and neuroinflammatory processes. We used miRWalk to predict genes relevant to neuronal processes that are putative targets of the three high-priority differentially regulated microRNAs. We filtered miRWalk predictions for those expressed in the DVC using available RNAseq data from rat models of autonomic dysfunction. Colored dots represent the target genes categorized based on gene ontology and functional annotation.

### Effects of persistent cardiac ischemia on transcriptional state

In order to investigate DMV homeostatic responses on a longer time scale, ligation of the proximal left anterior descending artery was performed with survival times of 1 week or 3 weeks to examine early dynamics of DMV responses to persistent ischemia. Similar approaches of assaying gene expression were used as with the RIPC studies. Through the use of a similar hierarchical clustering algorithm with average linkage of Pearson correlations, it is possible to identify 6 neuronal states and 5 gene clusters (GC) through dendritic tree height cut-offs that parsimoniously segregate groups. This provides the framework for understanding the heterogeneous responses to cardiac ischemia over time (Fig. 5a). The distribution of samples from experimental groups among these 6 SC is not uniform as would be expected if LAD ligation had limited effect on the behavior of the DMV as a whole. Instead, a clear increase in the representation of certain states with specific conditions and in many cases, distinct distributions of phenotypes on the left and right is observed (Fig. 5b). The 1 week sham (Sham-1) samples are generally divided into three main states, State-U, State-V, and State-Z. The 1 week LAD ligation samples (LAD-1) are distributed similarly, but with the additional representation in State-W. The sham 3 week samples (Sham-3) are represented in all SC with negligible left/right differences. The 3 week LAD samples (LAD-3) distribute much differently than all of the other groups; State-U and State-Z have only sparse representation whereas State-W, State-X, and State-Y are heavily represented, an effect that is exaggerated on the right side when compared to the left. Details of the gene order in the heatmap and sample composition can be found in Supplemental Table 2.

**Fig. 5:**
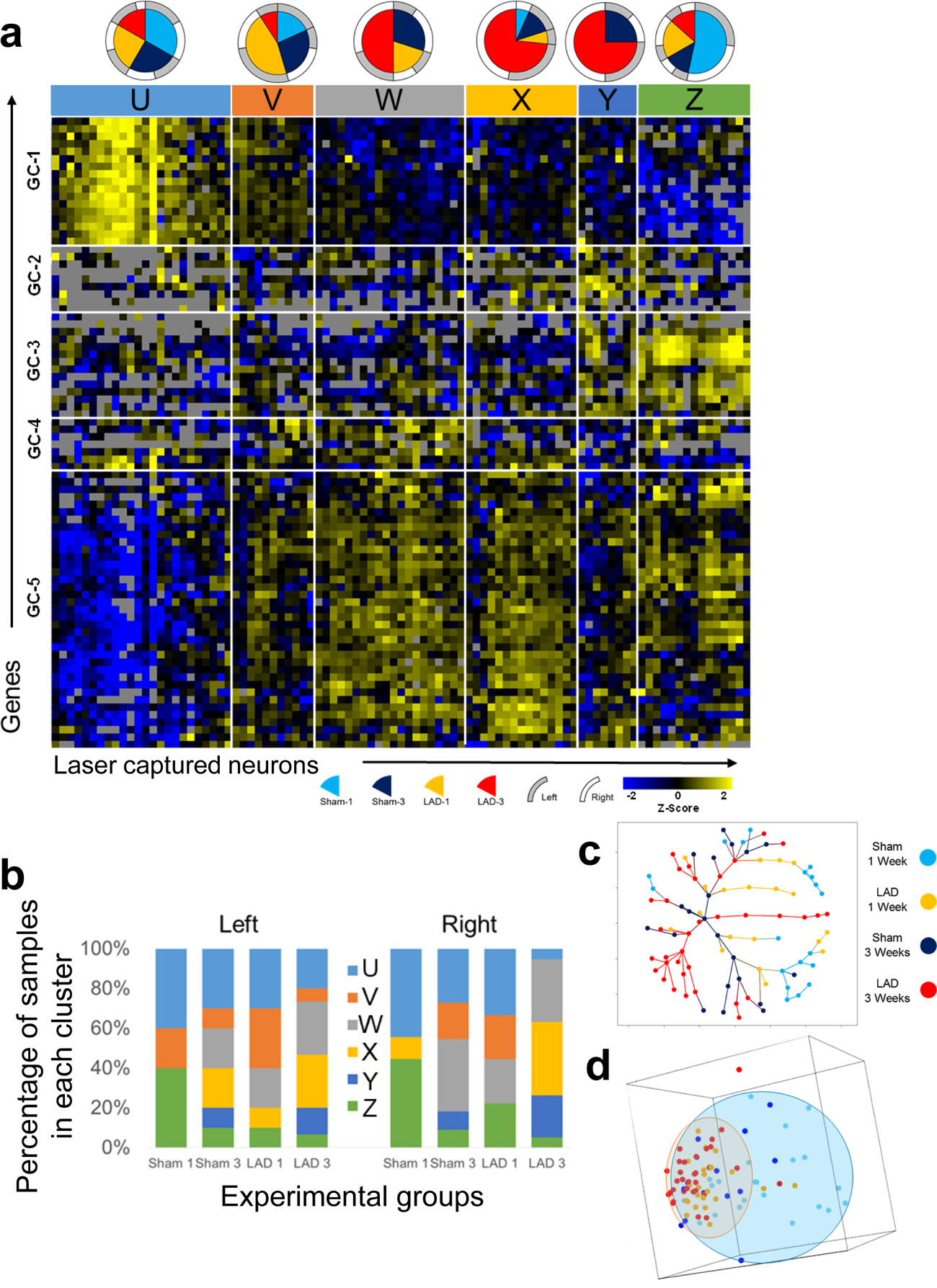
Chronic cardiac ischemia shifts the neuronal states of DMN neurons with distinct distributions at 1 week and 3 weeks. **a** Heatmap showing gene expression (Z-score of -ΔC_t_) organized by hierarchical clustering of both genes and samples as well as composition of sample cluster (pie charts). Each column represents one sample (pool of ∼5 neurons collected from only one side of the DMV). Each row represents a gene assayed through multiplex RT-qPCR. Sample clusters are colored for clarity of reference in subsequent Fig.s. Gray boxes indicate that no data is available for that particular reaction. **c,d** Dimensionality reduction through minimum spanning tree (c) and principal component analysis (d) demonstrates diminished expression space for LAD-3 samples toward the SC-C, SC-D, and SC-E expression patterns. Minimum spanning tree colored for experimental groups, shows the unique terminal branches of LAD-3 composed of samples of the SC-D and SC-E phenotypes. The same MST colored for neuronal state is shown in Supplemental Fig. S13. **d** Plot of first three principal components with ellipsoids showing the general expression space for the sham samples (blue) and the LAD samples (orange) regardless of time point. Points are colored according to the experimental group. This plot demonstrates a reduction in the expression space (diversity) of LAD samples toward an expression pattern that is present in some of the sham samples.

The distinguishing features of State-U are high expression of GC-1 and low expression of GC-5 with average levels of the others. State-V may be considered as a transition from State-U to State-W with expression levels of nearly all of the GC in between those of State-U and State-W. State-W has a characteristic high expression of GC-4 and GC-5 together and lower expression of GC-1 and GC-2. State-X has a high expression of GC-2 and GC-5 with more modest upregulation of GC-4 and downregulation of GC-3. State-Y includes upregulation of GC-2, GC-3, and GC-4 together. State-Z has a marked downregulation of GC-1 and marked upregulation of GC-3 with modest upregulation of GC-5. While not exact opposites, the composition of State-U and State-Z have gene clusters (GC-1, GC-3, and GC-5) that have opposing expression tendencies.

The two sample clusters at the opposite ends of the heatmap in Fig. 5a, State-U and State-Z, are the most clearly defined distinct transcriptional phenotypes based upon their average Euclidean distances in the minimum spanning tree (Fig. 5c, S13). Of note as well is that both State-U and State-Z are composed of samples from all of the experimental groups, albeit in different proportions, perhaps suggesting their ubiquity in DMV function, irrespective of cardiovascular perturbation. Clusters State-V, State-W, State-X may initially all appear to be transitional groups that represent the populations of neurons that are in the process of shifting from State-U to State-Z or vice versa. However, an examination of the minimum spanning tree (MST) in Fig. 5c suggests that State-V and State-W are more likely transitional clusters whereas State-X and State-Y are terminal phenotypes. The branching pattern for State-E in the MST suggests that it is really two separate phenotypes that in this case differ due to ischemic cardiac damage. There is one branch that is composed entirely of LAD-3 samples and has a distance of 3-4 in the MST from the bulk of the two main phenotypes, State-U or State-Z (Fig. S13). The other State-Y samples are from the Sham-3 group and are adjacent to the rest of the samples from State-Z.

A more broad interpretation of the main neuronal phenotypes considering all genes examined here suggests that State-U and State-Z are the phenotypes of DMV neurons that exist either at resting state or are recruited for generalized injury/stress response. This is shown in the distribution of sample clusters within experimental groups in Fig. 6b. The combination of nearly equal representation of all experimental groups for each and the distinct gene expression patterns support this. While the heatmap makes State-W and State-X appear very close in expression patterns, examination of the spanning tree suggests that State-X is more of a terminal state whereas State-W represents the transitional state. Given the poorly defined separation of these clusters, it is possible that they are a similar state, but are behaving in different ways, perhaps a result of inputs or feedback, but are operating under the same overall coordinated program. The main difference between State-W and State-X is the higher expression of GC-2 in State-X, a gene module that contains neuropeptides like somatostatin and galanin along with the somatostatin-3 receptor and the oxytocin receptor. It also has a clear upregulation of the beta-1 adrenergic receptor, better known for its cardiac specificity in the periphery, and for which little is known about function in neurons of vagal motor neurons. Another distinction includes the upregulation of GC-4 in State-W compared with State-X. GC-4 contains several inhibitory receptors: two GABA receptor subunits (ε and θ), two G_i_ coupled muscarinic receptors, and the G_i_ coupled alpha 2 adrenergic receptor. Coupled with the relative increased expression of several calcium channel genes save for Cacna1c (Supplemental Fig. S15), this is suggestive of a neuronal phenotype capable of pacemaker or even burst firing^25–28^. Also contributory to this conjecture is the upregulation of Grin2a, Hcn2, and Kcnn4, all of which are involved in autonomous firing^26, 29, 30^. Along with the unique inhibitory receptors from GC-4 there is also increased expression of Gabra1 and Gabrb1 GABA receptor subunits, which may be necessary to lower the resting membrane potential of these neurons or prevent large depolarization events to permit their function as autonomous pacemakers capable of burst firing^25, 26, 28^. Also of interest in State-W and State-X is the increased expression of several receptors that are responsive to two somatostatin receptor subtypes and both orexin receptor subtypes. This may suggest an increased interaction strength with the central nucleus of the amygdala and lateral hypothalamus respectively as the sources of somatostatin and orexins as neurotransmitters (discussed in supplemental text).

**Fig. 6.**
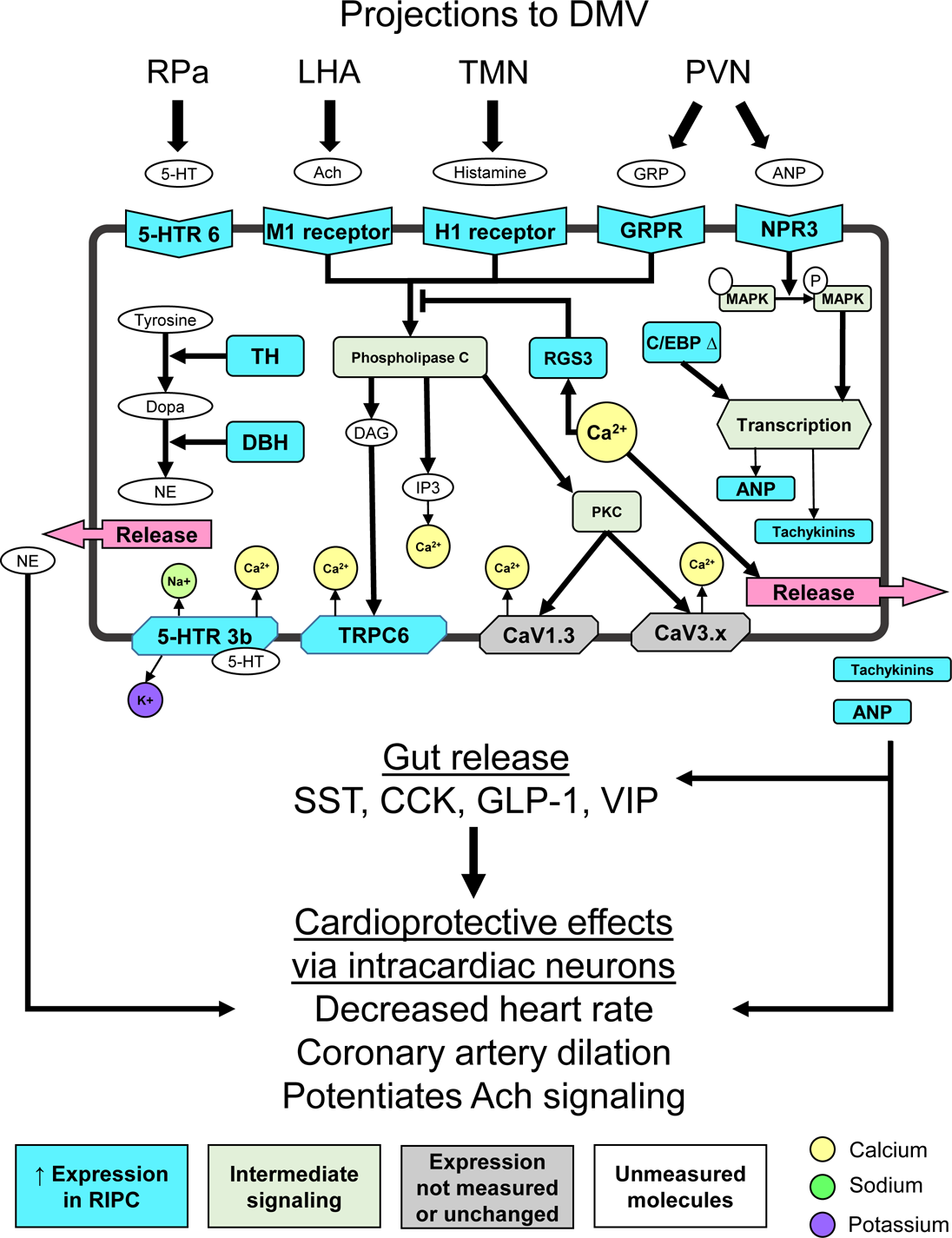
Plausible mechanism meditating RIPC effect in H1-receptor expressing DMV neurons. Schematic of H1-receptor expressing neuron highlighting potential mechanism based upon differential gene expression of receptors, intracellular effectors, and neuropeptides. Genes that are uniquely expressed in the H1 expressing neurons of the IPC cohort compared with non-H1 expressing neurons from the IPC group and compared with the H1 neurons of the naïve and sham cohorts. Brain regions likely acting on the receptors shown are shown along the top above the brackets. All pathway information was derived from literature inferences with several pathway connections representing multiple steps and intermediaries. The milieu of genes expressed suggest an increase in calcium influx that Gq signaling that drive the production and release of norepinephrine, atrial natriuretic peptide, and tachykinins from DMV efferent neurons to contribute to cardioprotection derived from remote, ischemic preconditioning (RIPC). Abbreviations: 5-HT 6 = serotonin receptor 6, M3 = muscarinic acetylcholine receptor 3, H1 = histamine receptor 1, GRPR = gastrin releasing peptide receptor, NPR3 = naturetic peptide receptor 3, MAPK = mitogen-activated protein kinase, TH = tyrosine hydroxylase, BH = dopamine beta hydroxylase, 5-HT 3b = serotonin receptor 3b, TRPC6 = transient receptor protein 6, DAG = diacylglycerol, IP3 = phosphotidyl inositol, Dopa = dopamine, NE = norepinephrine, 5-HT = serotonin, Ach = acetylchoine, GRP = gastrin releasing peptide, ANP = atrial naturetic peptide, Rpa = raphe pallidus, LHA = lateral hypothalamus, TMN = tuberomammillary nucleus, PVN = paraventricular nucleus.

Consideration of the expression pattern across all measured genes using principal component analysis (Fig. 6D) yields two interesting observations. The first is that the expression space, diversity of expression pattern, for the LAD ligation samples occupies a much more constrained space (red ellipsoid) than the sham samples (blue ellipsoid). The second observation of note is that this constrained space shown by nearly all LAD samples from both time points lies within the expression space defined by the sham samples. This suggests that there is programmatic coordination in response to ischemic injury and that this response is within the realm of normal expression patterns at least for some neurons. In this case the response phenotypes include State-V and State-W for LAD-1 and State-W, State-X, and State-Y for LAD-3.

In the consideration of gene expression correlation networks, the LAD-3 group has much lower connectivity than either of the 1 week groups and also less unique edges. Most surprising is the emergence of Pax4a as a unique hub gene in its relationship to many GC-5 genes. These networks also reveal an early response to LAD at one week that is uniquely and significantly coordinated as evidenced by the increased connectivity of the gene expression networks between the oppositely expressed GC-1 and GC-5. More detailed explanations, implications, an figues can be found in the Supplemental Text and Fig. 16-19.

### Effects of persistent cardiac ischemia on gene co-expression network topology

Gene co-expression networks were generated for each of the experimental groups using Pearson correlations and filtered using a q value cut-off of q<10^-3^. The unique edges for each of the networks generated are given in Fig. S16 with the connectivity in any of the experimental groups were excluded from the full network for each group given in Fig. S16-19. Genes with no connectivity in any of the networks based upon the cut-off criteria mentioned previously were excluded from all the network figues. It is clear from these networks that the LAD-1 group has the greatest overall connectivity with over half of all edges being unique that group. Many of these unique edges are the negative correlation between genes in GC-1 and GC-5. Also of note are unique edges of interconnectivity within GC-1 and GC-5 respectively. The notable unique edges of the Sham 1 week group include several negative correlations with Foxo4, Camk2, and Drd4 in GC-3 with genes in GC-1. Also, there are several unique positive correlations to genes in GC-5 from other GCs as well as some unique interconnectivity within GC-5. The Sham-3 group is most notable for its very sparse connectivity.

Within each of the treatment cohorts, there are several highly connected genes as is the baseline expectation for a biological network, suspected to have scale-free topology^31^. Overall, a few genes are highly connected across most of the cohorts, most notable being Cacna1b that is a member of the top five connected genes in each cohort. Cebpd is highly connected in the LAD-1 group and the LAD-3 group, but with many unique edges in the LAD-1 group. This suggests a role of Cepbd in mediating gene expression in the acute response to cardiovascular injury that persists to some extent beyond the acute phase.

### Putative DMV neuromodulatory network mechanism mediating RIPC-induced cardioprotective effects

While it is more difficult to elucidate a single differentiating factor from the chronic ischemia model, the RIPC model As one means of synthesizing these data, there is a significantly larger proportion of neurons expressing Hrh1 in the RIPC group, which begs the question whether this is due to a novel H1 expressing phenotype or if there are just more neurons of a similar phenotype. There are several genes that are differentially represented or expressed only in the H1 expressing neurons of the RIPC group. Based on the receptors expressed, these neurons appear to have the chance for increased influence from PVN, TMN, RPa, and LAH as compared with all other neurons from RIPC or other groups. A proposed mechanism is described in Fig. 6.

## Discussion

In this work, we aim to uncover the molecular substrates that underlie the DMV orchestration of the cardioprotective effects, providing insight as to what signaling cascades might mediate the effect as well as how the DMV itself integrates these signals and modulates target neurons in the gut, heart, and possibly other organ systems to promote cardioprotection. In order to do this, we first characterized the heterogeneity of DMV neurons at baseline, particularly with regard to their integrative input signal processing capacity leading to combinatorial neuropeptide and neurotransmitter production. Shifts in the distribution of these states occur dynamically in response to surgical stress and RIPC. Unique responses to three weeks of chronic cardiac ischemia include changes in gene expression and subsequent gene regulatory networks show a dramatic shift to a neurosecretory, enteroendocrine-like phenotype in a highly coordinated fashion. The dynamic changes in the DMV neuronal landscape appear to generally occur through selective recruitment of one state from others. The implications of these findings emphasize a need to consider the functional implications of neuronal heterogeneity with the understanding that at times only a subset of neurons from a brain nucleus may exert significant effects on physiology. These subsets may not be appreciated unless experimental work is done with sufficient granularity and time-dependent sampling.

The DMV is one of the few brain regions uniquely positioned to coordinate homeostatic organ function within the body. The anatomy of afferent connections and distribution of vagal efferents has been well known and this work provides the context needed to demonstrate the heterogeneity of DMV processing functionality. The distinct neuronal states that shift in adaptive or maladaptive directions provide a conceptual framework for thinking about how the DMV attempts to maintain the body’s homeostasis. Conditions like RIPC or other ischemic preconditioning may function as hormetic stresses that encourage the state shifts in the DMV necessary to manage the systemic effects of cardiac ischemia whereas the shifts observed after several weeks of persistent cardiac ischemia may demonstrate a compensatory response to augment the stress response.

In RIPC, there was a significant upregulation of the generally sparsely expressed H1 receptor gene (Hrh1), both in expression level as well as in number samples with expression above the limit of detection (Fig. S10b, S10d, S11). There is one known source of histaminergic projections to the DMV, the tuberomammillary nucleus (TMN) of the hypothalamus ^32–34^. Activation of the H1 receptor in the dorsal vagal complex has been shown to mimic the effects of satiety, reducing hyperphagia in rodent models of obesity ^35^. This is in line with its general ability to mediate depolarization in the low number of neurons in the DMV that express it ^32^. Such satiety signaling may be co-opted to mediate cardioprotective effects ^7, 23, 36^. While there is evidence that the TMN is responsive to cardiovascular perturbations and can indirectly modulate cardiovascular function ^37, 38^, this is the first indication we are aware of for its role in mediating cardioprotection in any sense. There are several genes that are differentially represented or expressed only in the H1 expressing neurons of the RIPC group. Of particular interest are significant decreases in expression in Sst and Cck. Somatostatin,delivered locally into the gut, decreases the release of somatostatin ^39^, cholecystokinin ^40^, glucagon-like peptide 1 ^41^, and vasoactive intestinal peptide ^42^, thus the decrease in Sst expression effectively indicates an increased chance for release of these peptides. Coupled with this decrease is a marked increased expression of Tac1 and Nppa. Neurokinins and atrial natriuretic peptide in the gut induce the release of the same four peptides mentioned above along with several others ^43–48^. Apart from action at the gut, both tachykinins and ANP have well-described effects on the heart including decreased heart rate ^49, 50^, coronary artery vasodilation ^51^, and potentiation of acetylcholine signaling ^52, 53^. Further, the combination of norepinephrine and ANP likely works on post ganglionic neurons in the heart to mediate sympathoinhitibiton that was once thought to require input from at least two different neurons ^54–56^, but we now show may plausibly come from the same vagal preganglionic neurons. There are several receptor and ion channels whose expression is increased and whose coordinated effects would lead to increased excitability and release of peptide transmitters ^57–61^. There is also a shift from high expression of Rgs2 in other neurons to higher Rgs3 expression. Rgs2 is an inhibitor of G_q_ and enhancer of G_i/o_ signaling whereas Rgs3 is responsive to increased levels of Ca^2+^ as a feedback regulator of G_q_ signaling, not a tonic inhibitor like Rgs2. Based on the receptors expressed, these neurons have the increased influence from PVN, TMN, RPa, and LAH as compared with all other neurons from RIPC or other groups. This suggests a novel circuit for centrally mediated cardioprotection that may be exploited to stimulate an effect similar to RIPC.

While a distinct subset of neurons mediate the effects of RIPC over the course of hours, there are significant shifts in the population landscape of neuronal states after three weeks of chronic cardiac ischemia. These shifts occur from the recruitment of neurons to a putative neurosecretory potential, particularly on the right side of the DMV more than the left, as shown in State-W, State-X, and State-Y. This programmatic response appears mediated, at least in part, by Pax4a, a transcription factor is strongly regulated by the transcriptional repressor REST which drives stem cells into a neurosecretory phenotype ^62, 63^. Disinhibition of gene expression through the downregulation of REST with concomitant upregulation of Pax4a is restricted largely to neurons, but also occurs in developing/regenerating pancreatic islet cells, in particular β and δ cells ^64^. The coordination of increased expression of the genes in GC-5 may be accomplished in part through effects of Pax4a as evidenced by its unique network connectivity in that condition (Supplemental Fig. S16). Coupled with the coordinated upregulation of Cacna1b, Cacna1d, Cck, Gabra1, and Gabra2 the pictures is painted less of a neuron and more of an enteroendocrine cell, like a pancreatic β/δ cells or colonic L/I cells ^65–67^. Canonical satiety peptides (CCK, GLP-1, SST, etc) have a protective effect on peripheral organs in the setting of ischemic damage at the heart ^36, 68–73^. That the DMV may deliver peptides directly to postganglionic neurons in the periphery adds dimension to the concept of vagal tone as a mediator of health and disease.

The National Institutes of Health (NIH) programmatic initiative aimed at development of means to Stimulate Peripheral Activity to Relieve Conditions (SPARC) was motivated by therapeutics derived from electrical stimulation of the vagus nerve in multiple contexts. We highlight here the complexity and plasticity of the vagal efferent neurons from the DMV. We further show that acute DMV responses only involve a subset of neurons mediated by distinct effector neurotransmitters and neuropeptides. This suggests there are molecular means by which stimulation of autonomic peripheral activity may also involve specific neuropeptides or other effectors to mediate more specific effects. As we continue to personalize medicine, we will continue to specify therapeutics. This work provides a framework for investigations to uncover more specific means by which the autonomic nervous system mediates physiological homeostasis and compensates for disease states.

## Methods

### Animals

This study used adult male Sprague Dawley rats in the weight range 250-300g (Envigo). All animal experiments were carried out in accordance with protocols approved by the Thomas Jefferson University Institutional Animal Care and Use Committee.

### Procedure for remote ischemic preconditioning (RIPC)

The remote ischemic preconditioning (RIPC) protocol used here largely mirrors that described by Gourine (Mastitskaya et al. 2012; Basalay et al. 2016) with all protocols being approved by the Thomas Jefferson University Institutional Animal Care and Use Committee. Adult 250-300g male Sprague Dawley rats (Envigo) were given a single intraperitoneal injection of ketamine/xylazine (100 mg/kg and 10 mg/kg) and then subjected to either a sham surgery or the RIPC protocol. Both surgeries involve bilateral small incision in the upper thigh, isolation of the femoral vein and artery from the nerve in the femoral sheath, and placement of a suture thread around only the vessels. In the RIPC procedure the thread was tied to occlude the vessels, being left untied in the sham, for fifteen minutes. The suture was removed and the limbs permitted to reperfuse. Both the ischemia and reperfusion were confirmed through visualization of blood flow and pallor followed by subsequent recoloration of the surrounding muscle tissue. The animals survived for two and half hours further before sacrifice. During this survival period, the animal was placed on an isothermal heating pad and the wounds in the lower limbs covered with gauze pads soaked in sterile saline.

### In vivo manipulations-ligation of the left anterior descending artery (LAD)

All experiments were performed with the approval of the Institutional Animal Care and Use Committee at Thomas Jefferson University. All procedures and sacrifices were done within the same four hour circadian period. Male Sprague-Dawley rats (250-300g) were anesthetized with ketamine (100mg/kg) and xylazine (10mg/kg) and intubated with an 18g venous catheter and the animal was ventilated with oxygen-enriched room air using a rodent ventilator (Harvard Apparatus). A left thoracotomy was performed using aseptic technique and the heart was exteriorized. The left anterior descending (LAD) artery ligation was performed by passing a 6-0 prolene cardiac suture around the LAD artery and tying it off. Paling of the myocardium was observed before the heart was replaced back into the thoracic cavity and the thoracotomy closed with 4-0 silk suture. The sham ligation procedure involved the same preparations and left thoracotomy, but instead involved passing the 6-0 suture and needle around the LAD artery without tying the suture. The animals were then extubated and allowed to recover on a heating pad for four hours before being transported back to the holding facility where they were maintained in a 12h light-dark cycle and given *ad libitum* access to food and water. Animals were monitored post-surgically every 24 hours for signs of distress or improper healing and removed from the study if any such signs presented.

### Animal Sacrifice and Harvesting of Brainstem

After the appropriate survival period (1 to 3 weeks for LAD/Sham experiments, 2.5 hours for RIPC experiments), each animal sacrificed by rapid decapitation that was preceded by brief 60 seconds of 5% isoflurane in O_2_. The brain was quickly removed and placed into ice cold artificial cerebrospinal fluid for separation of the brain stem that was then rapidly frozen in optimal cutting temperature (OCT) medium for cryosectioning. The heart was removed and rinsed in cold PBS before being rapidly frozen in OCT. No more than ten minutes passed between decapitations and freezing the tissue in OCT for each animal.

### Processing of Samples for LCM

The brain stems were sectioned at 10μm and collected on SuperFrost Plus glass slides in preparation for laser capture microdissection (LCM). Neurons in the DMV were identified by use of a rapid (∼15 minutes) immunofluorescence labeling procedure staining for NeuN. Single neurons and pools (composed of 2-5 neurons) were collected from the DMV of adult male Sprague Dawley rats (N=3 animals) in the region coextensive with the area postrema using the Acturus HS Capsure system. Each cap containing neurons was visualized after capture to ensure that only neurons were included in the samples. Any cap containing non-neuronal cells were not used in the study. Apart from collecting neurons only, some samples were collected from the same slides that included the entire area of the DMV represented on that slice. The samples for this study were all collected from the part of the DMV that begins at the obex and runs caudally for 500μm.

### Rationale for RT-qPCR

Morphological studies on DMV neurons have suggested for some time that there is some heterogeneity within the population^74, 75^. Such work has led to uncovering of multipolar peripherally projecting efferent neurons, multipolar GABAergic interneurons, and a population of unipolar neurons ^18^. It has also been shown that there some organization within the nucleus as to the efferent vagal targets with interneurons interspersed ^75^. To date, there has been significant work attempting to identify the molecular biology that underlies DMV function, but through examination of only a handful of transmitters or peptides at a time. Many of the co-labeling studies that were performed found some populations of dual labeled cells, suggesting that co-expression of essential neurotransmitters or receptors is more the rule than the exception, a concept that will be further developed in the work that follows. This work aims to understand the molecular heterogeneity of DMV neurons through broad examination of gene expression. In order to reliably assay gene expression at the level of single cells, it was decided to take a targeted multiplex RT-qPCR approach rather than RNA-seq. In the majority of single cell RNA-seq experiments, it is very difficult to detect low levels of expression in a reliable and repeatable way, often requiring reliance on the gold standard of PCR to confirm essential findings. Further, it is often the case that a cohort of a few hundred genes are found to be differentially expressed and even less than that to find biological relevance. Our approach considers a targeted set of 100-200 genes that are highly likely to have functional implications to DMV effector function, based upon the body of published literature accumulated to date on DMV in a variety of contexts and disease states (Supplemental File 1). Until the sensitivity and specificity of RNA-seq for single cells or larger scale targeted gene expression techniques like MER-FISH are refined, any approach will inevitably miss something, whether through technical limitations or incomplete target gene selection. Our approach here provides data with very high fidelity from which analysis and reasoning can be carried forth with confidence.

### Selection of genes for RT-qPCR

The multiplex RT-qPCR experimental design included genes, which are listed in Supplemental Tables 1 and 2. The selection of these genes from the thousands expressed in the DMV was accomplished through inference based on our previous work in the dorsal vagal complex and through analysis of publicly available transcriptomic studies. There is, however, one dataset examining the DMV in human patients with Parkinson’s disease (PD) that may be helpful in understanding DMV pathology (GDS4154). Through use of ANOVA and template-matching, several genes that were differentially expressed in PD patients and only changed in the DMV were considered in constructing the gene list for this current work. Further justification for gene assay selection can be found in the Supplemental File 1.

### High throughput RT-qPCR

The respective sets of genes (Supplemental Tables 1 and 2) were measured in each sample using multiplex RT-qPCR via the Biomark system from Fluidigm, as described previously ^12^. These samples were randomly allocated to three different microfluidic chip multiple qPCR runs along with samples across treatment groups to minimize technical noise across sample types. For naive animals 84 samples passed quality control composed of 60 single cell samples and 24 cell pools totalling 156 cells in all. 106 samples passed quality control comprised of 45 RIPC and 61 sham from N=3 animals for each cohort. Pooled single cell samples were prepared for multiplex real-time quantitative polymerase chain reaction (RT-qPCR) with the Biomark HD system using VILO III (ThermoFisher) reverse transcription of RNA into cDNA followed by 22 cycles of pre-amplification (Taqman Pre-amp Master Mix) of the cDNA using the collection of primers that were utilized in the multiplex RT-qPCR. Each set of primers was designed to span introns when possible and was validated through *in silico* PCR (Primer BLAST, NCBI) and observation of a single band with the expected amplicon size on PCR gel using standard rat brain RNA, prepared for PCR by the same methods outlined previously ^12^. The quality of the results of the multiplex RT-qPCR was ensured through examination of the melt curves and standard curves. If there were multiple peaks for the melt curves or if a reliable standard curve was not present, the assay was not used in further downstream analysis. Similarly, if a sample showed that more than half of the assays were below the limit of detection, the entire sample was not utilized in downstream analysis. Similar techniques were utilized for the chronic ischemia experiments with the gene assays and numbers enumerated in Supplemental Table 2.

### Normalization and Handling of Missing Information

Raw C_t_ values from the multiplex RT-qPCR that passed melt curve based quality control were first median centered within each sample in order to account for the variations in total RNA in the sample to get delta C_t_ values. For analysis and reporting purposes, negative delta C_t_ (-dC_t_) values were used because a lower C_t_ value is representative of a higher expression level. By using -dC_t_ values, higher values now represent higher gene expression and are normalized to account for different initial total RNA amounts. One difficulty with assaying single cells in general is the large number of assay-sample combinations that end up having values below the limit of detection, resulting in a large number of null or not applicable (NA) values. Most of the algorithms used to cluster across several measurements omit NA values or impute them. Omission of NA values may lead to an over-estimation of expression for a given group or cohort due to not counting NA values as being of very low to zero expression. They are instead considered to be missing information. In this case, the most likely reason for an NA value is very low to no expression of a certain gene which is different from uncertainty about its expression. Similarly, imputing NA values when there is such a large number will likely introduce patterns that are overfit from the non-NA data. In order to work around this, a technique of binning the data was utilized solely for the purpose of determining membership in clusters/phenotypes. Five bins were used with the lowest being composed of all NA values and the remaining four being the quartiles of the remaining non-NA values. This method permits the clustering algorithms to use the information from a lack of expression in the calculations and generate a more accurate membership in a cluster for each sample accordingly. For the reporting of data and any associated statistics, expression values in the form of negative delta Ct values (centered on the sample median of the top 35 expressed genes) are used along with the percentage of samples in a given group that have expression values above the limit of detection. For bar plots, expression values are recentered as a whole such that no value is below zero, making them visually easier to interpret. This also effectively makes zero the limit of detection and the scale of log2 relative expression where a difference of 1 indicates a two-fold difference in expression value and a difference of 2 indicates a 4-fold difference, and so on.

In order to statistically compare groups, it was necessary to consider both the expression value measures of central tendency (means and medians) as well as the number of samples that had a value above the limit of detection. As mentioned previously, the NA values that normally are ascribed to data points where there is uncertainty in the expression value due to technical problems or very low expression appeared to come primarily from very low expression in this set of experiments. This can be seen in the presence of primer dimers in the qPCR melt curve analysis in samples that otherwise have clear signal in more than 25% of all assays. The presence of primer dimers suggest that the reaction chemistry is proceeding appropriately, but there is insufficient template to amplify. When samples that had globally low signal across all assays were removed, the NA values that remain are far more likely to result from being below the limit of detection rather than technical problems. It is this line of reasoning that led to the use of NA values as data the analyses conducted here. It should be noted that a high number of NA values are not generally handled well in dimensionality reduction, even with imputation and ascribing an actual value to the NA based upon a fixed distance below the lowest -ΔCt value introduced a regularity in the data that doesn’t scale accordingly with the potential for washing out the signal.

For purposes of heat map visualization, the negative delta Ct values (normalized by sample to adjust for starting RNA content through centering on median gene expression value), can then be centered or normalized within each gene. Normalization may be done through calculation of a Z score for each value within a gene, giving the number of standard deviations from the mean (e.g. Z=1 is one standard deviation above the mean and Z=-2 is two standard deviations below the mean). This was the approach taken to visualize the data in Fig. 6. As mentioned previously, all statistics were performed on the negative delta Ct values, not the gene centered or normalized values.

### Other Statistical Analysis

In order to find statistical differences between treatments and over time in the chronic ischemia model work, a mixed linear model ANOVA, aggregating the pooled single cell samples up to the animal level, was used with Tukey post-hoc analysis to correct for multiple comparisons. For determinations of gene expression correlation, each sample was taken to be independent and pairwise Pearson correlations between genes were computed. All statistical calculations were performed using R statistical software.

### Prediction and Testing of microRNA regulators of DMV Gene Expression in RIPC

Given the large cohort of genes that had decreased expression in the RIPC cohort, microRNAs emerge as a potential regulator. Although microRNAs might act on gene expression in a myriad of ways, searching only the 3’UTR of genes increases the chance that the microRNA that binds there plays an inhibitory role in expression of the mRNA target ^76^. Through combined use of four algorithms, we determined the cohort of microRNAs that most parsimoniously lead to decreased expression, a cohort of 20 genes that were decreased in RIPC. If a microRNA had a putative binding site in the 3’UTR in more than one algorithm, it is shown in Fig. 5A. The algorithms used here were miRWalk ^77^, RNA22 ^78^, Targetscan ^79, 80^, and miRanda ^80^.

Broad assays of mircoRNA were obtained through the examination of differentially expressed microRNAs in the DMV of male rats. In this study, four conditions were tested: LAD alone, rIPC alone, rIPC followed by LAD two hours later, and age-matched control. Animals were sacrificed and brains were harvested 24 hours after LAD in the case of LAD alone and rIPC followed by LAD and 26 hours after rIPC alone. Animals were sacrificed and brainstems were harvested as described above and were sectioned at 200 μm and the DVC was extracted using a tissue punch 1mm in diameter. Samples were collected and assayed for microRNA content using Nanostring as previously described ^81^. After normalization and quality control, 146 microRNAs remained with detectable expression. Pavlidivis Template Matching was used to find microRNAs whose expression was perturbed in LAD alone compared to AMC, but whose expression returned to the AMC baseline with rIPC preceding LAD. Current literature on these three microRNAs suggest many interactions with ion channels and other excitatory and neuronal processes, supporting our prediction that perturbation of these microRNAs could provide some level of protection to reduce cardiac pathology and malfunction following LAD.

## Data Availability

The authors declare that all the data supporting the findings of this study are available within the article and its supplementary information files or from the corresponding author upon reasonable request. All the raw and processed HT-qPCR data as well as Nanostring microRNA profiling data is available online in the GEO database under SuperSeries GSE155053 which contains links to the individual data sets (HT-qPCR data: GSE155047 and GSE155049; Nanostring data: GSE154990).

## Supporting information

Assay Selection Literature

## Acknowledgements

Financial support for this work was provided by the National Institutes of Health under National Heart, Lung and Blood Institute grant U01 HL133360 and under the Stimulating Peripheral Activity to Relieve Conditions (SPARC) program grant OT2 OD023848 (PI: Kalyanam Shivkumar) subaward to J.S.S. and R.V.

## Author Information

### Affiliations

Daniel Baugh Institute for Functional Genomics and Computational Biology, Department of Pathology, Anatomy and Cell Biology, Thomas Jefferson University, Philadelphia, PA Jonathan Gorky, Alison Moss, Marina Balycheva, Rajanikanth Vadigepalli & James S. Schwaber

## Author Contributions

J.G., R.V., and J.S.S. conceived and designed the study. J.G. conducted the experiments and high-throughput qPCR assays and analyzed the single neuron gene expression data. M.B. performed the surgeries and cardiac histopathology analysis. A.M. performed nanostring microRNA profiling and analyzed the data. J.G. drafted the manuscript with revisions from A.M., R.V., and J.S.S. J.S.S. supervised the study.

## Ethics Declarations

Competing interests: The authors declare no competing interests.

## SUPPLEMENTAL TABLES AND FIGURES

**Supplemental Fig. S1:**
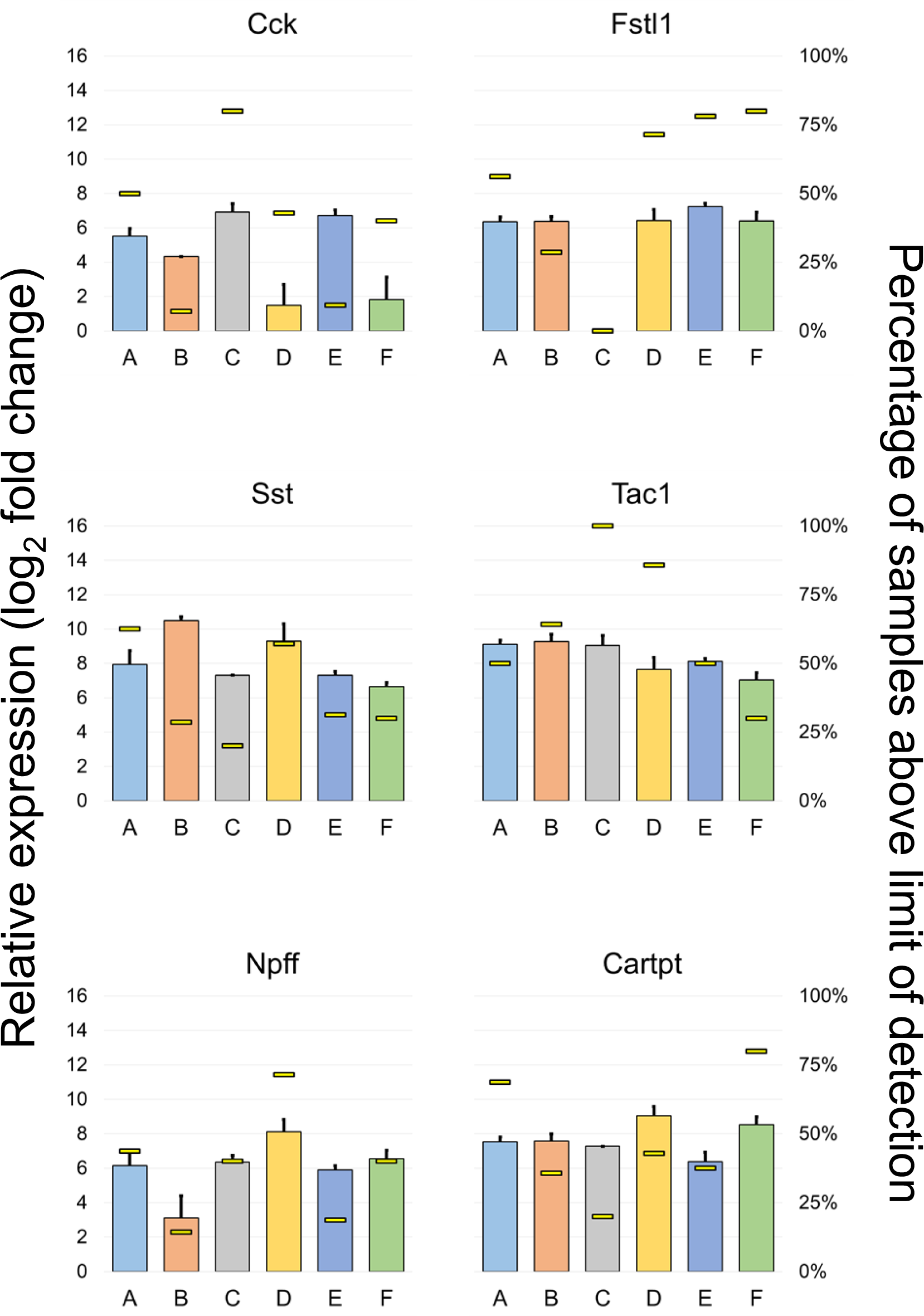
Sample cluster expression of genes that may indicate multiple peptidergic states. As with other figures, the left axis corresponds with the colored box with error bar (SEM) that represents gene expression (log2 relative gene expression with zero as limit of detection). The right axis represents the percentage of samples within each cluster that have an expression value above the limit of detection and is shown by the narrow bar in each group.

**Supplemental Fig. S2:**
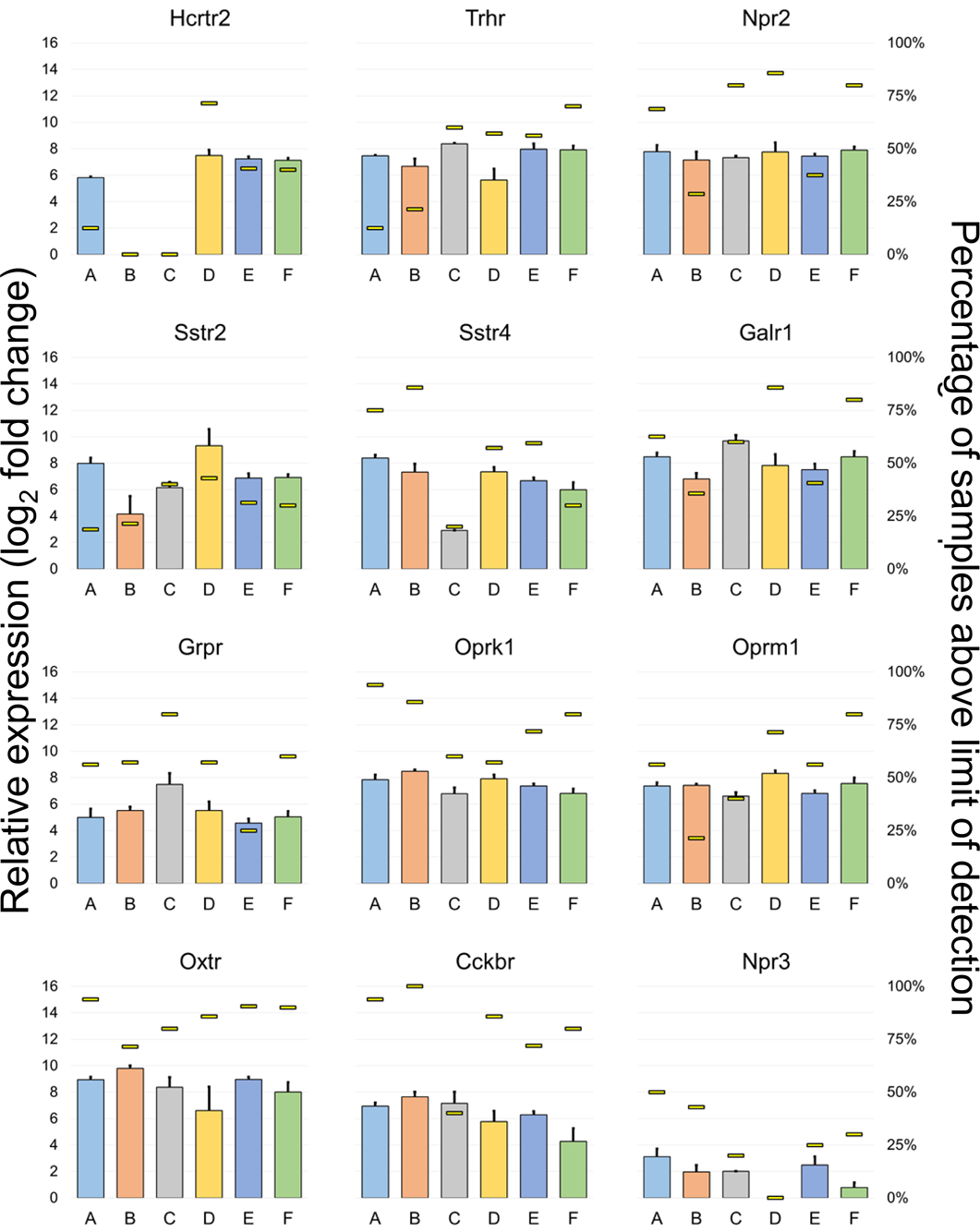
Sample cluster expression of genes that code for peptide receptors may help determine which neurons receive peptidergic inputs from specific regions of the brain. As with other Fig.s, the left axis corresponds with the colored box with error bar (SEM) that represents gene expression (log2 relative gene expression with zero as limit of detection). The right axis represents the percentage of samples within each cluster that have an expression value above the limit of detection and is shown by the narrow bar in each group.

**Supplemental Fig. S3:**
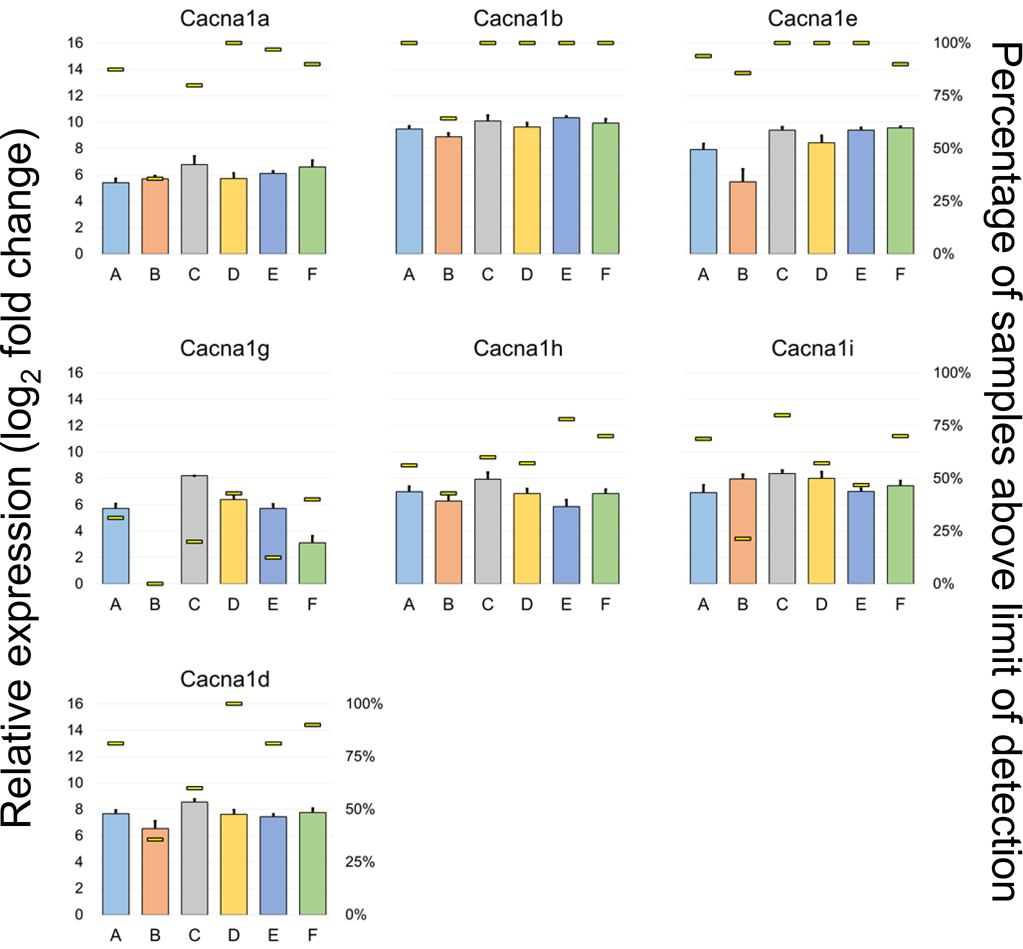
Sample cluster expression of genes that code for calcium channel subunits. Due to the consideration of calcium fluxes in DMV neurons driving apoptosis as described by Surmeier along with the generation of pacemaker-like activity (Goldberg et al. 2012), the distinct expression of calcium channel mix is of interest here. While there is general consistency in the high-voltage gated calcium channel subunits (Cacna1a, Cacna1b, and Cacna1e), there is a great deal of heterogeneity among the low-voltage gated channels (Cacna1g, Cacna1h, Cacna1i). This may suggest differences in burst firing or peacemaking activity. As with other figures, the left axis corresponds with the colored box with error bar (SEM) that represents gene expression (log2 relative gene expression with zero as limit of detection). The right axis represents the percentage of samples within each cluster that have an expression value above the limit of detection and is shown by the narrow bar in each group.

**Supplemental Fig. 4:**
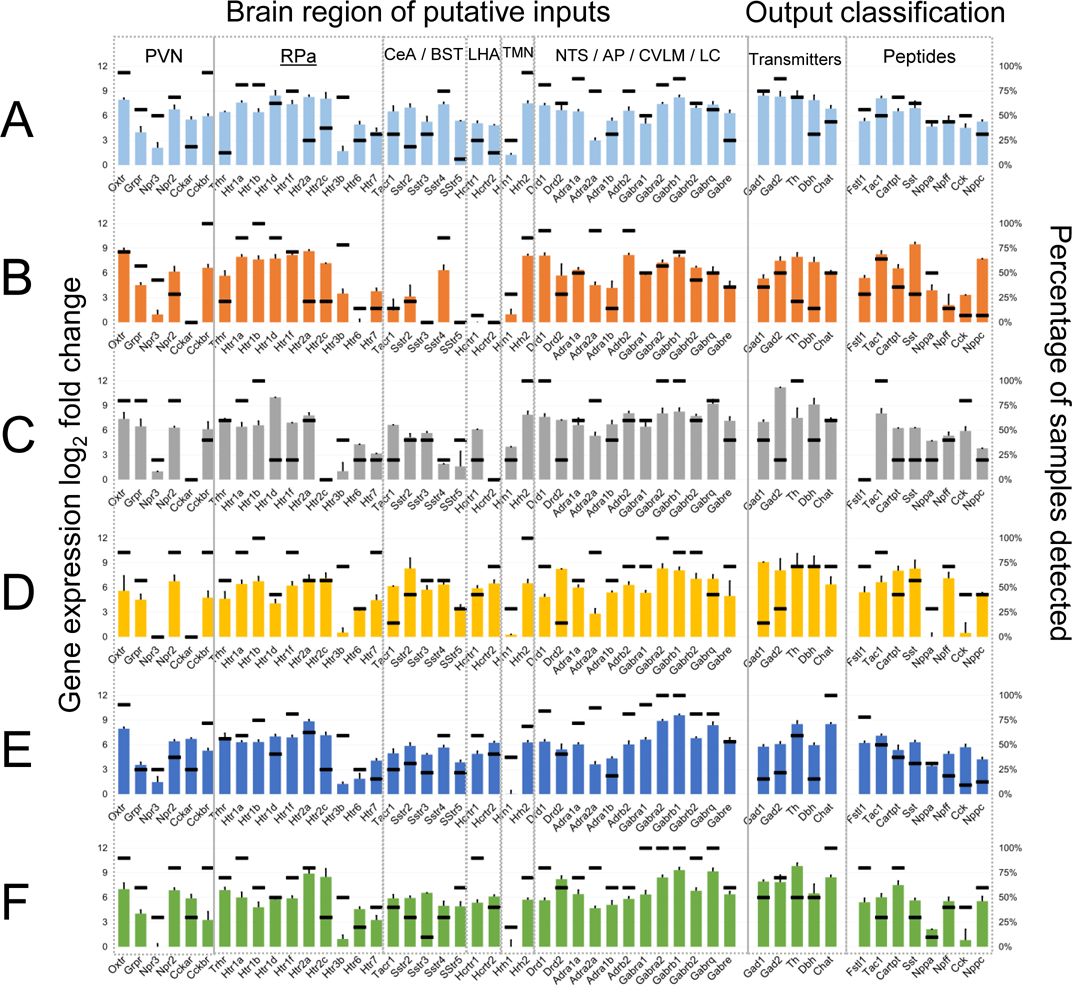
Datasets showing gene expression and sample proportion data for each state. These particular data were used to construct the input/output maps as shown in Fig. 2.

**Supplemental Fig. S5:**
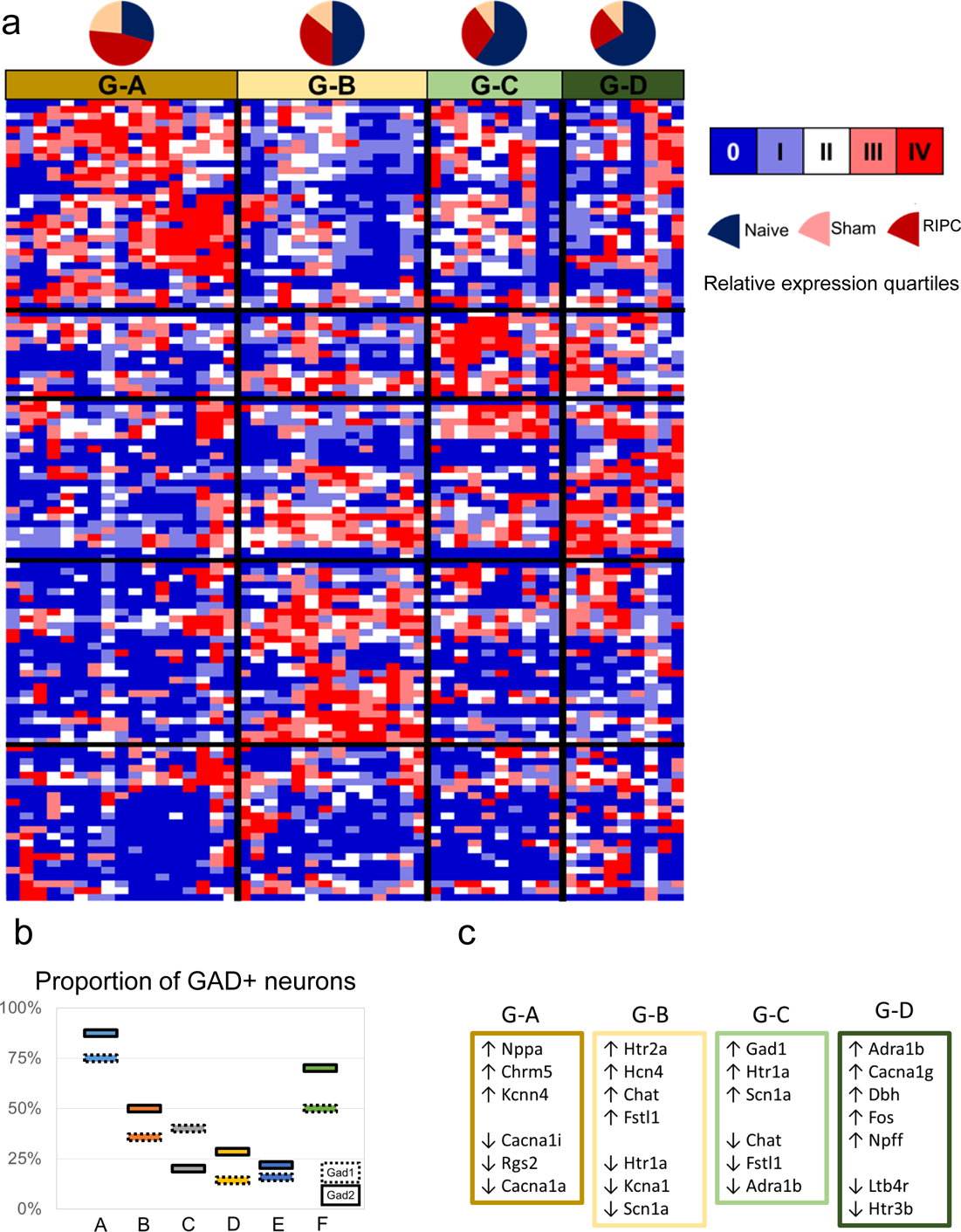
Heatmap of Gad+ single cells with distributions from experimental treatment groups. Pooled samples were omitted from this aspect of the analysis in order that anything found to be co-expressed with Gad1 or Gad2 could be attributed to a single cell as real co-expression. Four clusters emerge from this analysis that were determined without regard for the clusters found in Fig. 3a in order that the most potentially meaningful and strong correlations could be identified.

**Supplemental Fig. S6:**
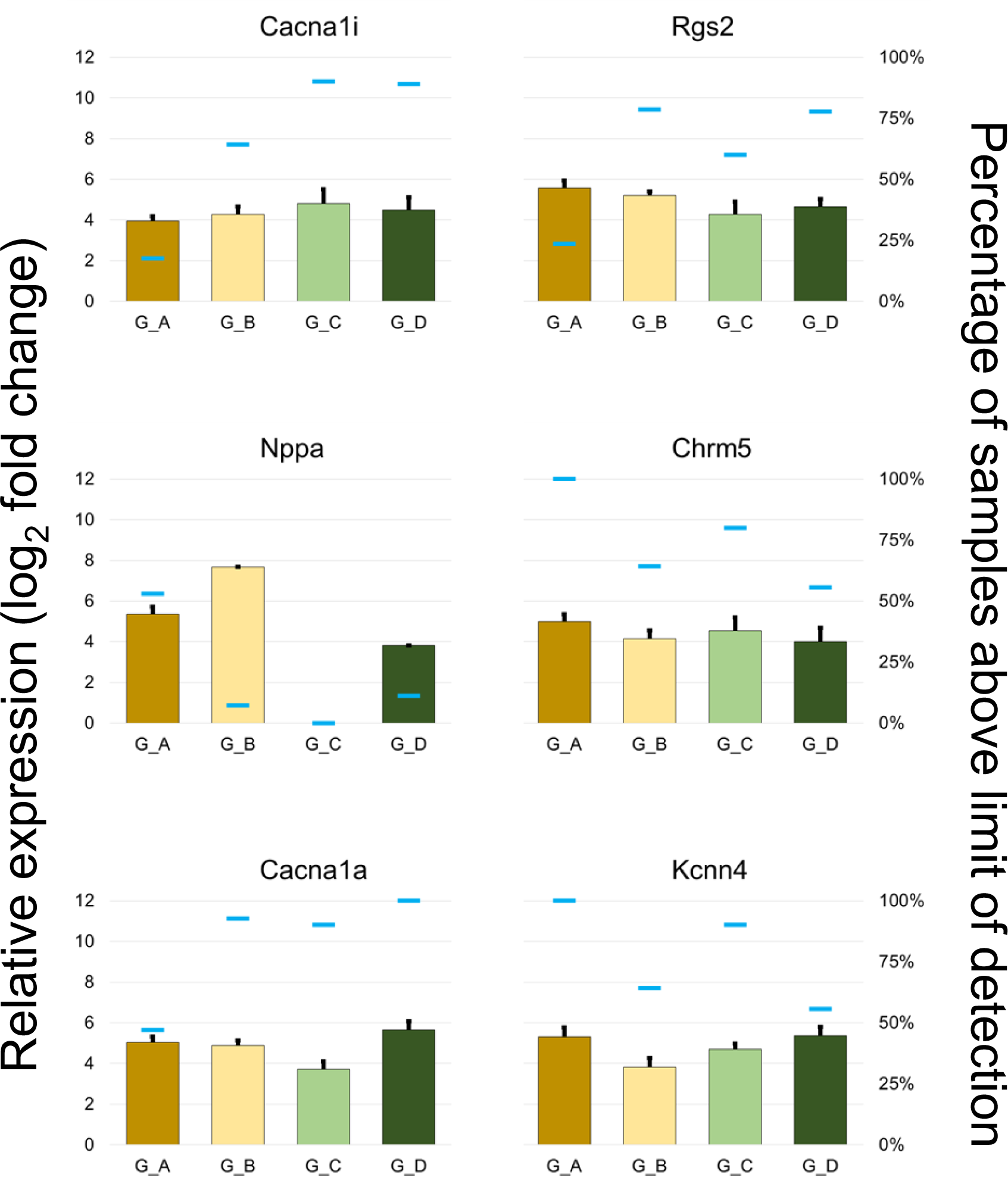
Genes with distinct expression patterns in cluster G_A. Binomial testing with a cutoff p<0.001 comparing G_A with at least two other groups was used to determine inclusion in this set. As with other Fig.s, the left axis corresponds with the colored box with error bar (SEM) that represents gene expression (log2 relative gene expression with zero as limit of detection). The right axis represents the percentage of samples within each cluster that have an expression value above the limit of detection and is shown by the narrow bar in each group.

**Supplemental Fig. S7:**
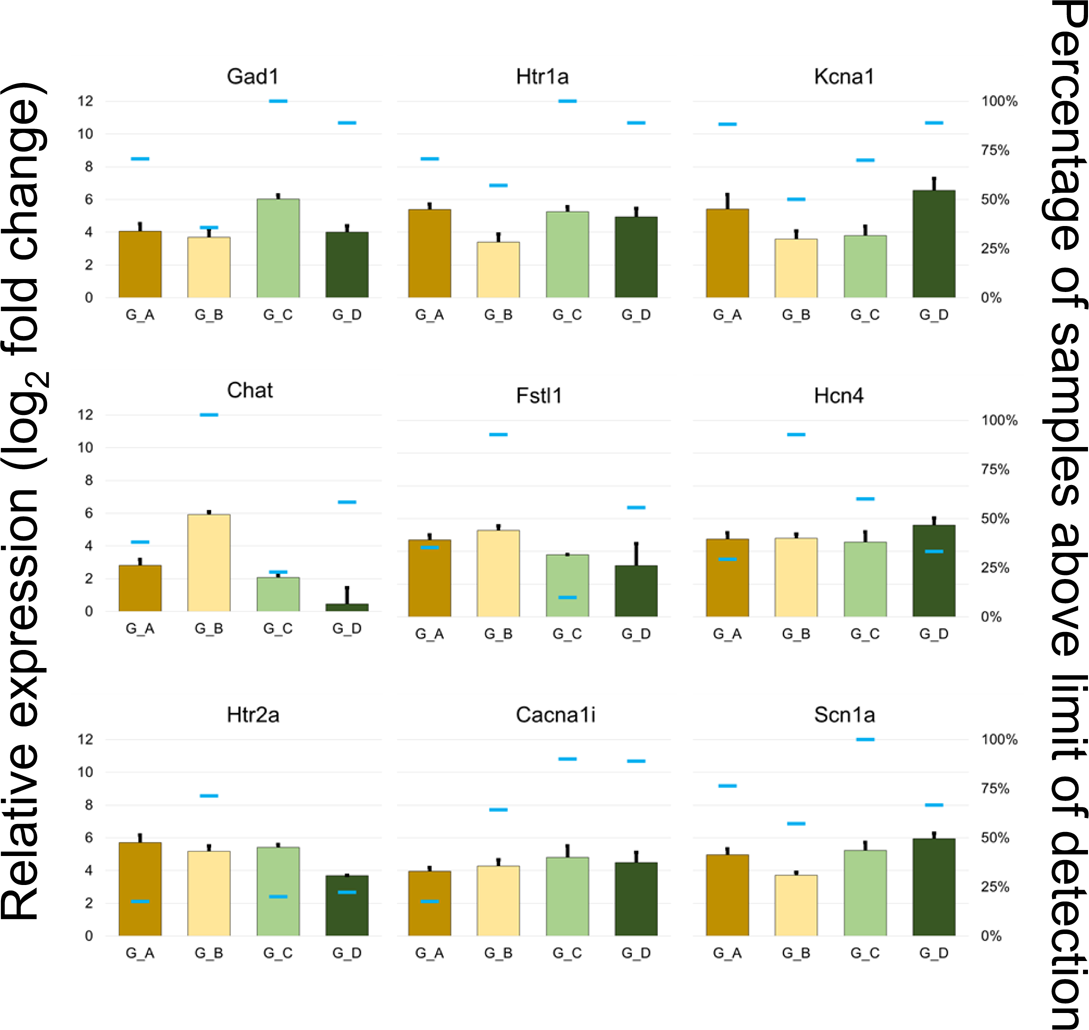
Genes with distinct expression patterns in cluster G_B and/or G_C. These clusters are included together due to the often opposite extremes in gene expression. Many of the genes collected here not only have significant differences in number of neurons with expression above the limit of detection, but also in the levels at which the gene is expressed in such neurons (ANOVA with Tukey post-hoc p<0.05, Exceptions: Hcn4, Htr2a, and Cacna1i). Binomial testing with a cutoff p<0.001 comparing G_B or G_C with at least two other groups was also used to determine inclusion in this set. As with other Fig.s, the left axis corresponds with the colored box with error bar (SEM) that represents gene expression (log2 relative gene expression with zero as limit of detection). The right axis represents the percentage of samples within each cluster that have an expression value above the limit of detection and is shown by the narrow bar in each group.

**Supplemental Fig. S8:**
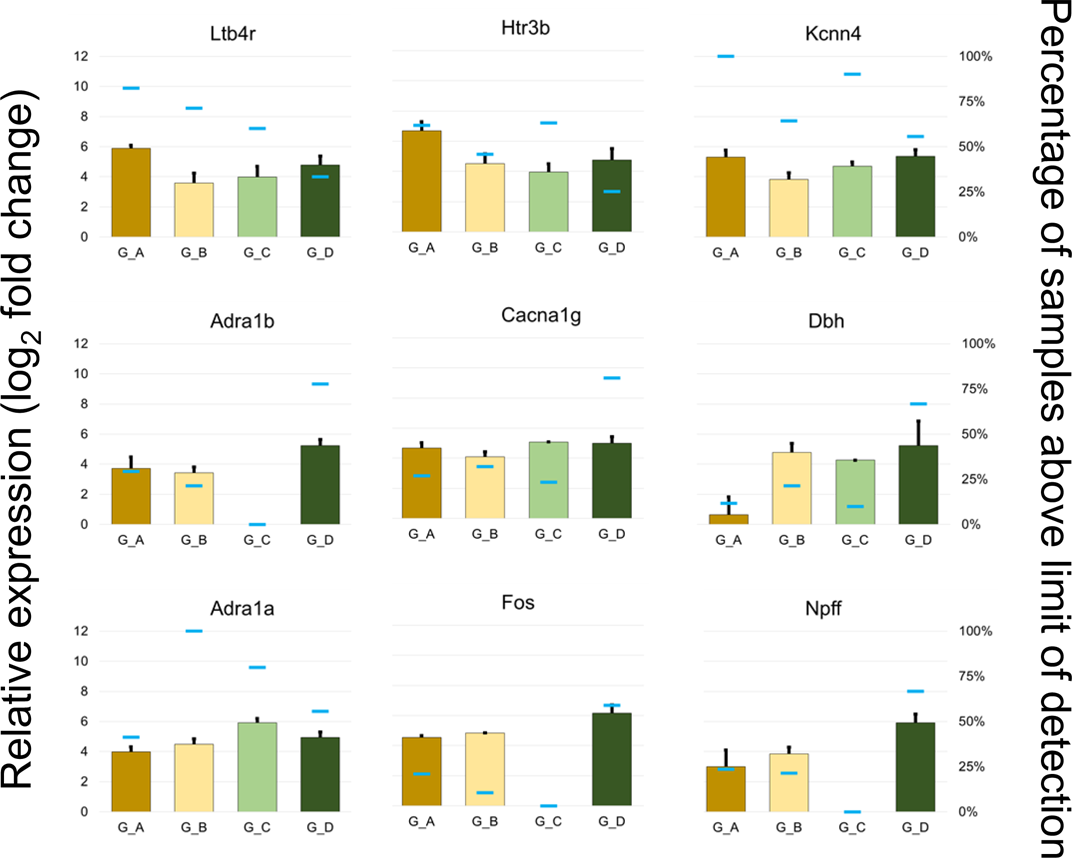
Genes with distinct expression patterns in cluster G_D. Binomial testing with a cutoff p<0.001 comparing G_D with at least two other groups was used to determine inclusion in this set. As with other Fig.s, the left axis corresponds with the colored box with error bar (SEM) that represents gene expression (log2 relative gene expression with zero as limit of detection). The right axis represents the percentage of samples within each cluster that have an expression value above the limit of detection and is shown by the narrow bar in each group.

**Supplemental Fig. S9:**
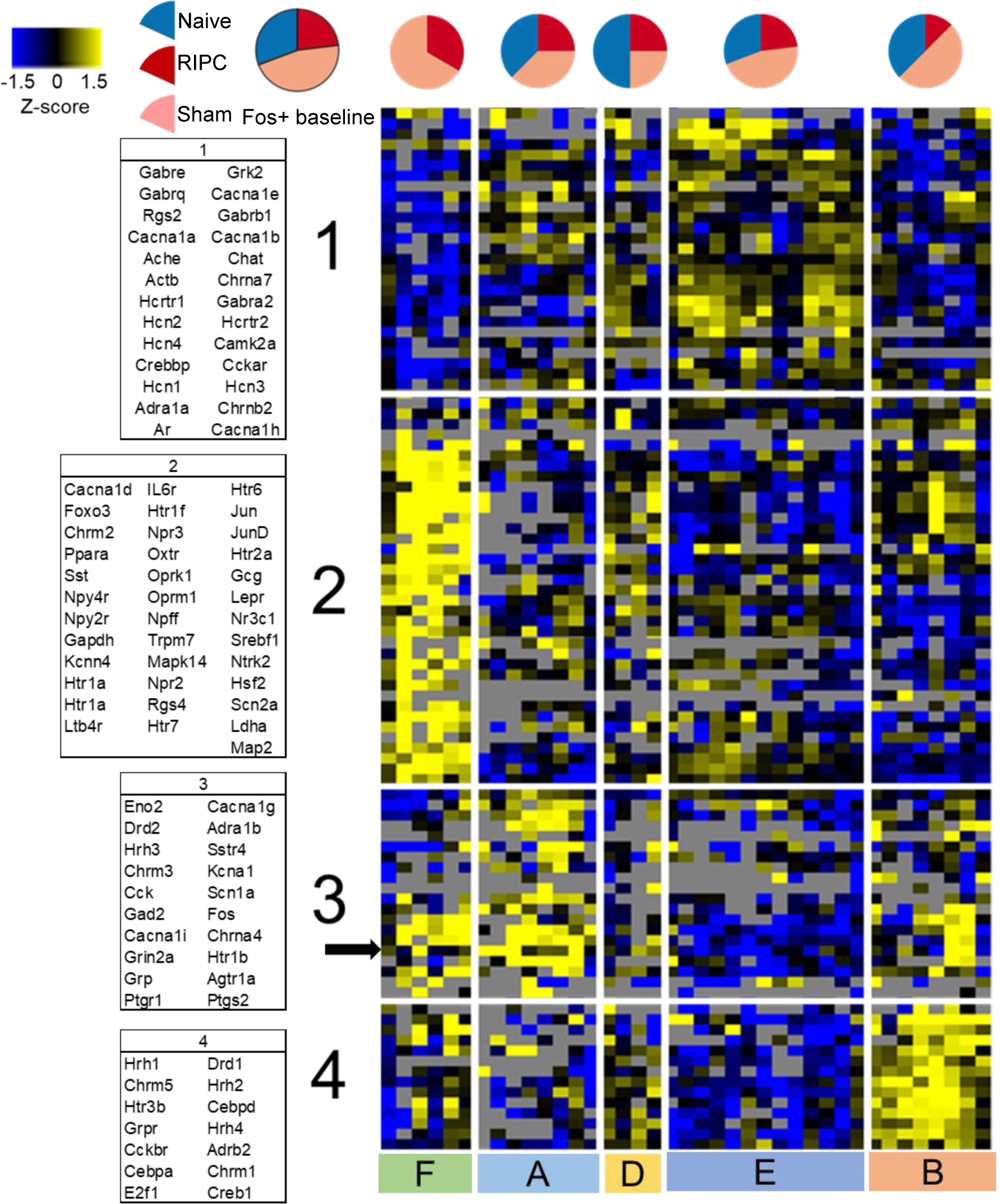
Heatmap showing only Fos+ samples suggests characterization of a surgical effect, but not a specific RIPC effect. Heatmap is colorized by Z-score (Z±1.5) within Fos+ samples only. Genes that comprise each cluster are given to the left of the heatmap, going down the columns then from left to right ascertain gene order on the heatmap. The row corresponding to Fos is shown by the black arrow in gene cluster 3. While the sample clusters of Fig. 1B were not completely recapitulated here, there are similar patterns that emerged from the *de novo* hierarchical clustering algorithm performed on this subset.

**Supplemental Fig. S10:**
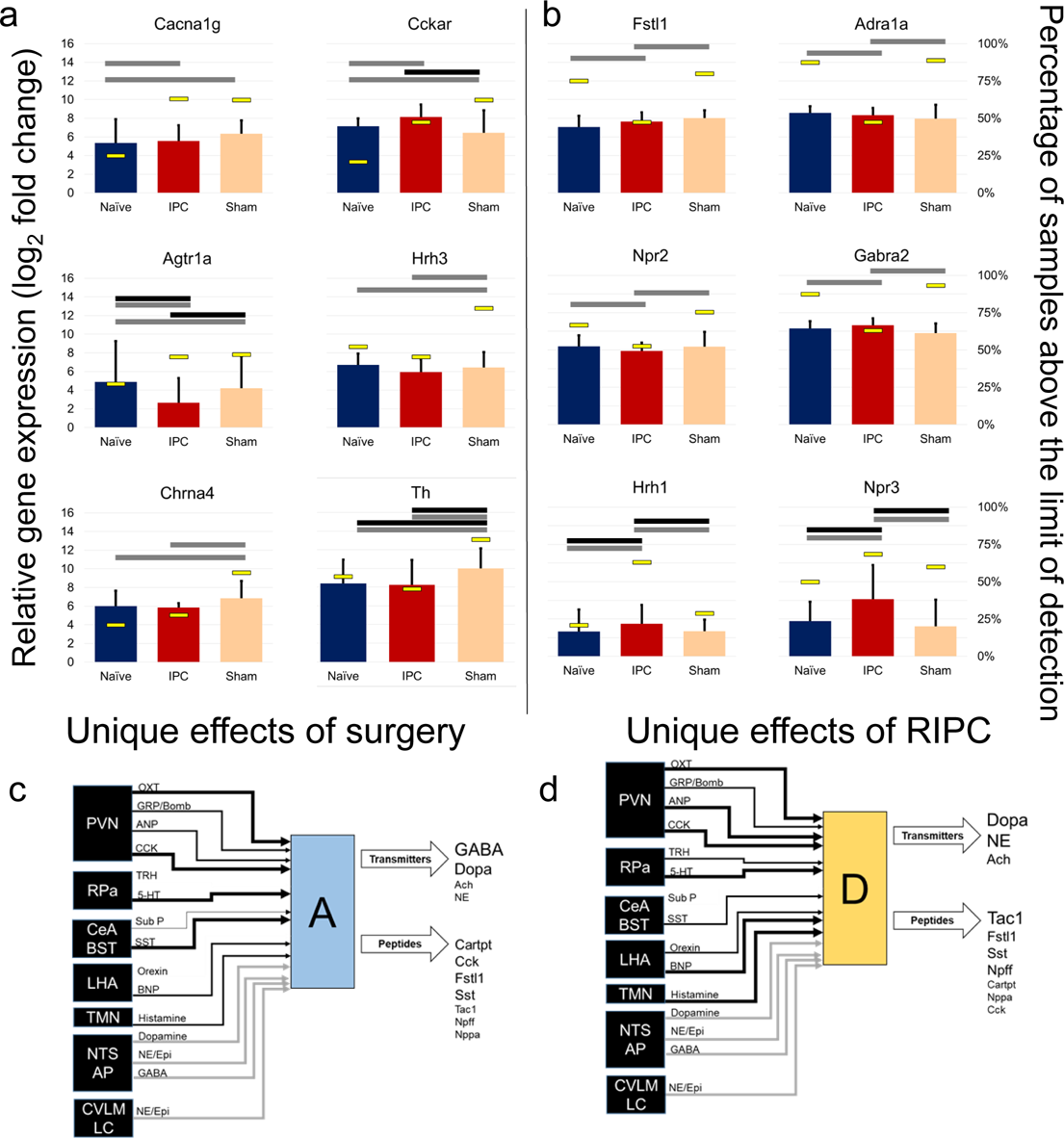
**a** In a demonstration of the effects of the stress of surgery or anesthetic (Sham), several genes were differentially expressed in both the RIPC and sham cohorts as compared with the naïve. In the case of Hrh3 and Chrna4, there appears to be a sham only effect, suggesting that the surgical treatment effects are mitigated by the addition of transient peripheral ischemia **b** Genes altered uniquely in RIPC versus either sham or naïve cohorts. Data shown here is for pooled samples only in order to generate the most accurate direct comparisons. For each of these genes, the single cell data mirrored the pooled sample data. The cut-off criteria used was a combination of binomial testing (gray bar, p<0.001) and/or pairwise T-test (black bar, p<0.05) to determine differences in percent of samples above limit of detection and expression values of expressing samples. Bars shown are mean±SEM. **c** Neuronal state A with disproportionate association with the Sham condition as shown in state A_2_. (d) Neuronal state D contains several uniquely upregulated genes in RIPC.

**Supplemental Fig. S11:**
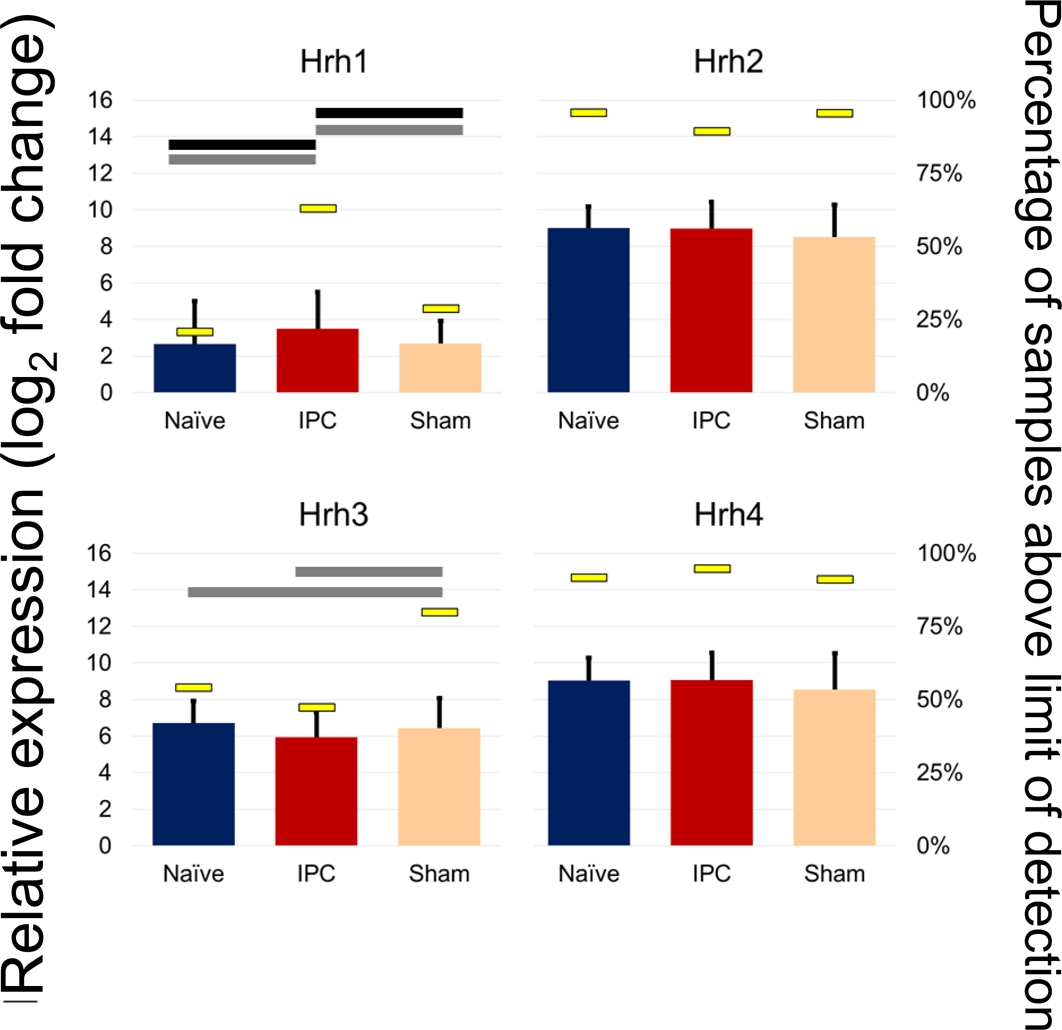
Histamine receptor subtype expression in response to RIPC and sham surgery. While the surgery itself leads to an increase in Hrh3 expression, this is suppressed in the RIPC to remain at naïve levels. In place of an increase in Hrh3 is an increase in Hrh1. The cut-off criteria used was a combination of binomial testing (gray bar, p<0.001) and/or pairwise T-test (black bar, p<0.05) to determine differences in percent of samples above limit of detection and expression values of expressing samples. As with other Fig.s, the left axis corresponds with the colored box with error bar (SEM) that represents gene expression (log2 relative gene expression with zero as limit of detection). The right axis represents the percentage of samples within each cluster that have an expression value above the limit of detection and is shown by the narrow bar in each group.

**Supplemental Fig. S12:**
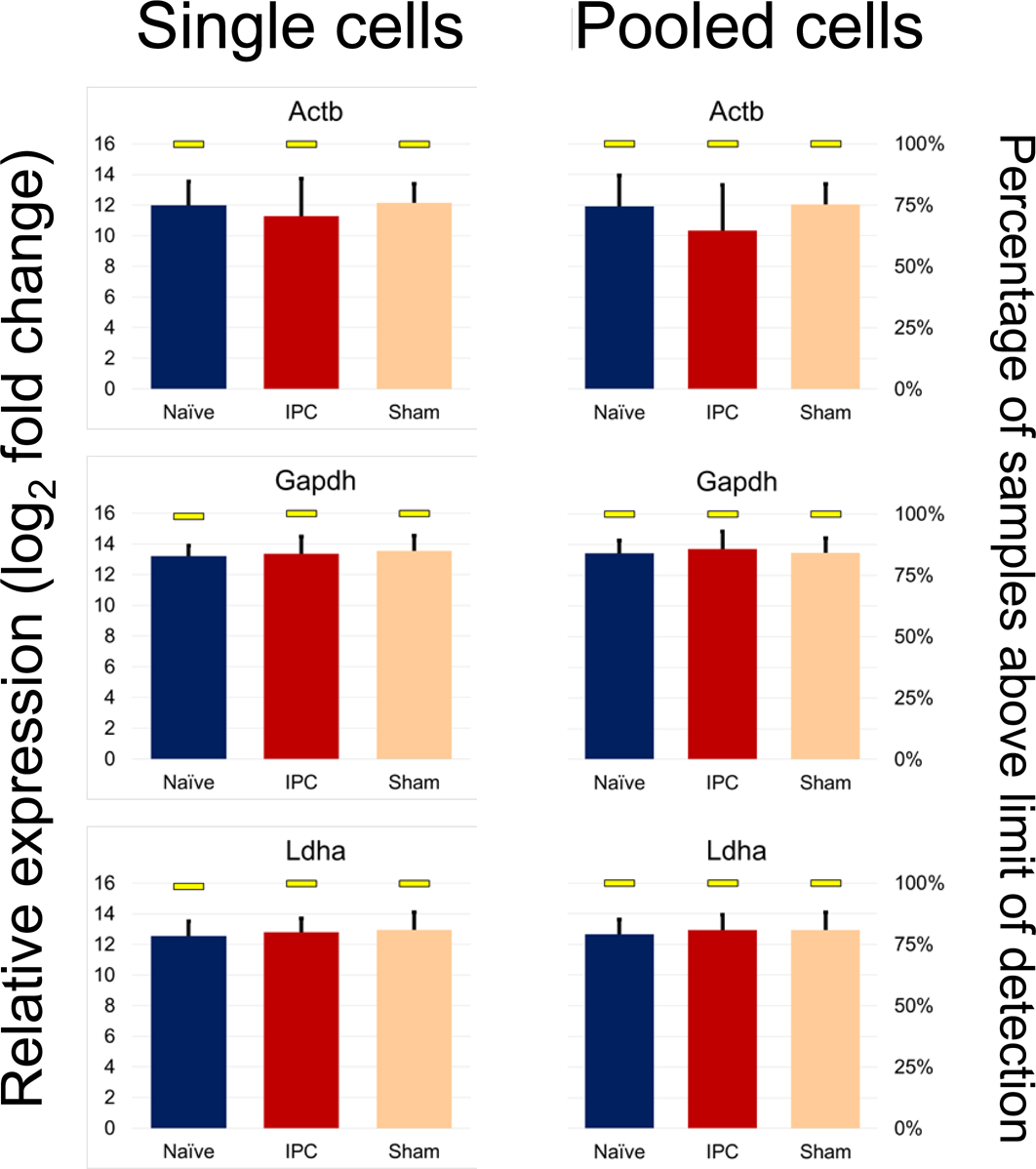
Reference genes show no differences between treatment groups nor between single cells and pooled single cell samples. Given the differing number of single cell and pooled samples within each treatment group at baseline, it is important that the technical differences in assaying single cells versus pools not persist after sample median centering normalization. As with other Fig.s, the left axis corresponds with the colored box with error bar (SEM) that represents gene expression (log2 relative gene expression with zero as limit of detection). The right axis represents the percentage of samples within each cluster that have an expression value above the limit of detection and is shown by the narrow bar in each group.

**Supplemental Fig. S13:**
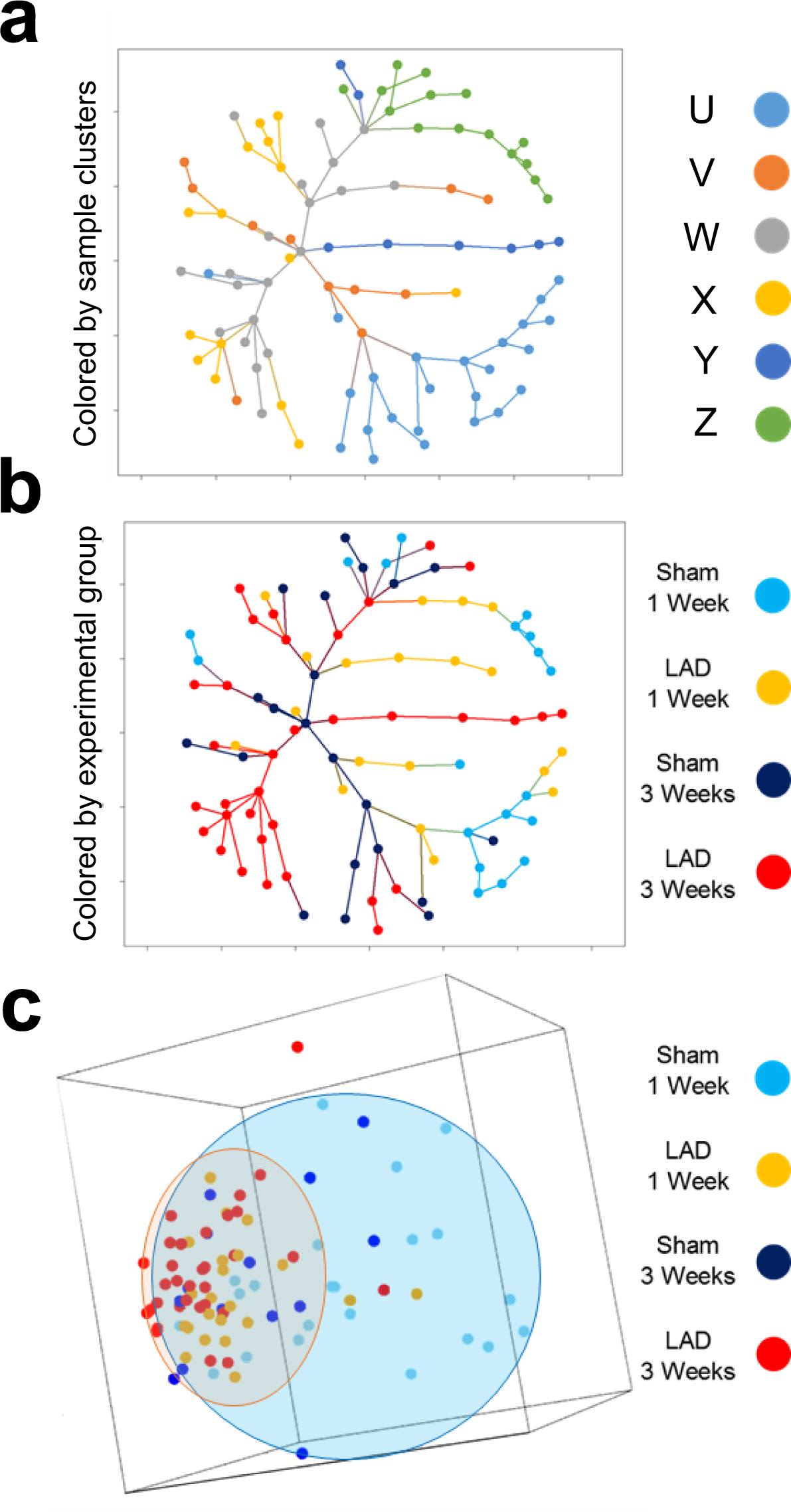
Dimensionality reduction though minimum spanning tree and principal component analysis demonstrates diminished expression space for LAD-3 samples toward the SC-C, SC-D, and SC-E expression patterns. **a** Minimum spanning tree colored by sample clusters shows that SC-B and SC-C are more transitional phenotypes with SC-A, SC-D, SC-E, and SC-F being more terminal. **b** The same minimum spanning tree as in **a**, but colored for experimental groups, shows the unique terminal branches of LAD-3 comprised of samples of the SC-D and SC-E phenotypes. **c** Plot of first three principal components with ellipsoids showing the general expression space for the sham samples (blue) and the LAD samples (orange) regardless of time point. Points are colored according to experimental group. This plot demonstrates a reduction in the expression space (diversity) of LAD samples toward an expression pattern that is present in some of the sham samples.

**Supplemental Fig. S14:**
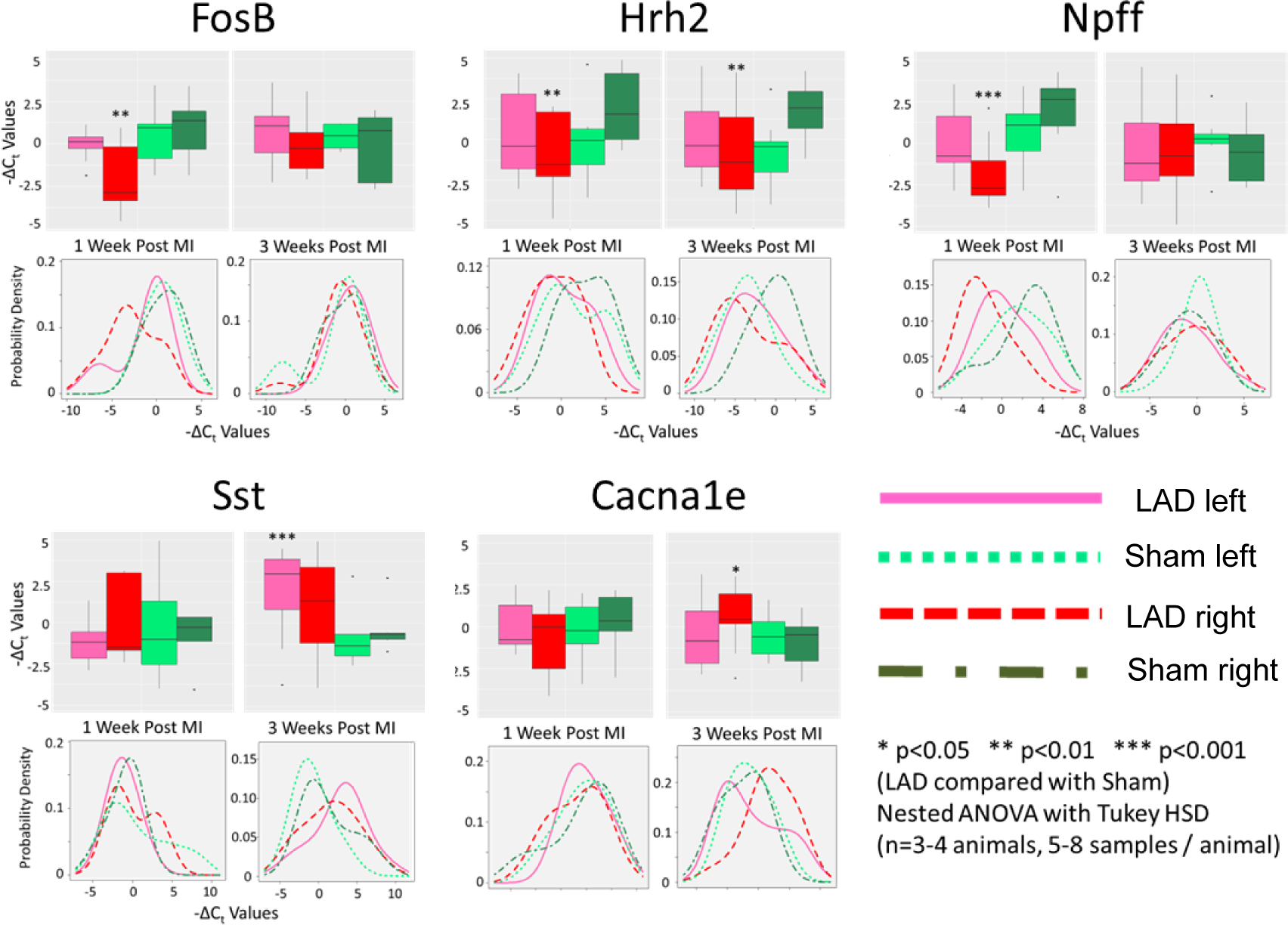
Genes with statistically different expression between experimental groups showed bilateral asymmetry. Of all the genes examined, only five showed statistically distinct expression levels between sham and LAD for either time point when accounting for animal level effects using a mixed linear model nested ANOVA (lme4 package in R). For each gene, boxplots of gene expression values are shown broken down by quartiles. Asterisks (*) indicate statistically significant differences between the sham and LAD surgery groups within a given time point on either the left or right side (as indicated by box/line color). Also shown for each gene is a density plot for the expression values for both left/right and sham/LAD conditions within a time point.

**Supplemental Fig. S15:**
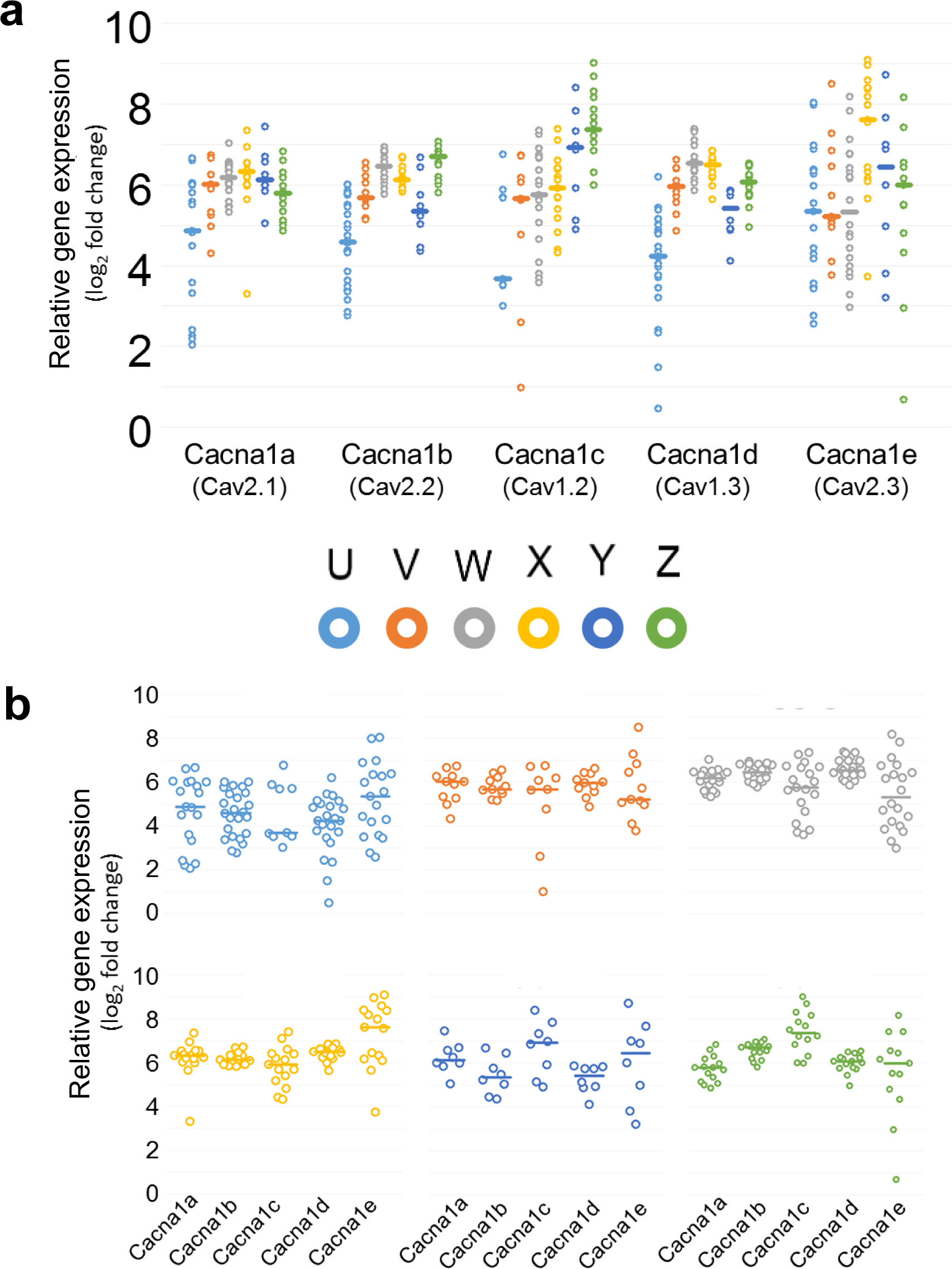
Expression of voltage gated calcium channels across samples clusters show some distinct patterns that may hint at functional behavior. **a** Each calcium channel’s gene expression values across all sample clusters with each dot representing one sample of pooled neurons. **b** Each sample cluster’s pattern of calcium channel expression with each dot representing a pooled neuron sample. Both **a** and **b** show the same data points, only grouped differently to aid in making comparisons.

**Supplementary Fig. S16:**
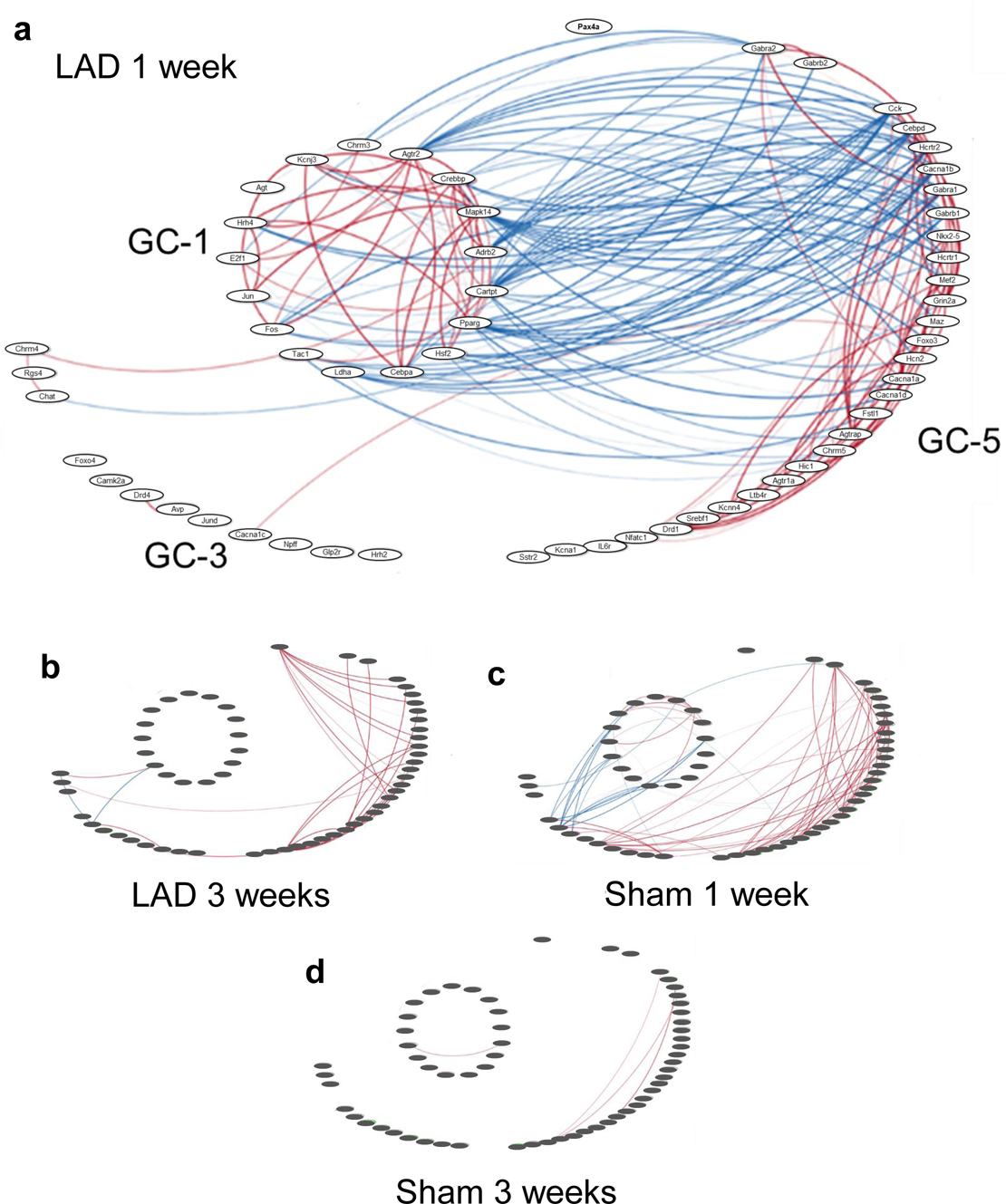
Correlation networks of each experimental group showing only edges that are unique to that group. Edges are derived from Pearson correlations with cutoff of q<10^-3^. Blue edges are negative correlations and red edges positive with the edge thickness proportional to the correlation coefficient. **a** Network for LAD-1 group and nodes enlarged to show gene names. **b** Network for LAD-3 group. **c** Network for Sham-1 group. **d** Network for Sham-3 group.

**Supplementary Fig. S17:**
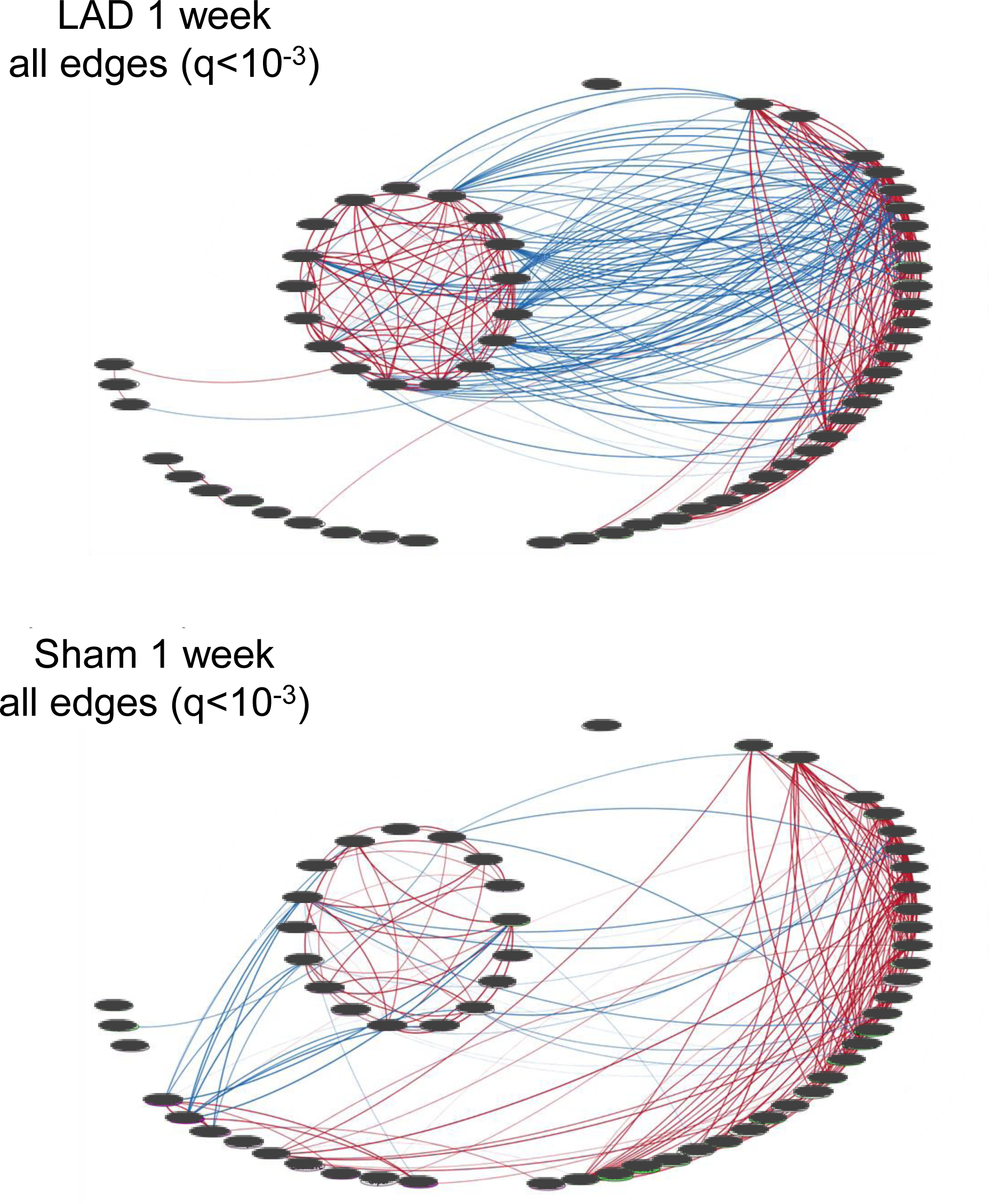
Correlation networks of each experimental group showing all edges. Edges are derived from Pearson correlations with cutoff of q<10^-3^. Blue edges are negative correlations and red edges positive with the edge thickness proportional to the correlation coefficient.

**Supplementary Fig. S18:**
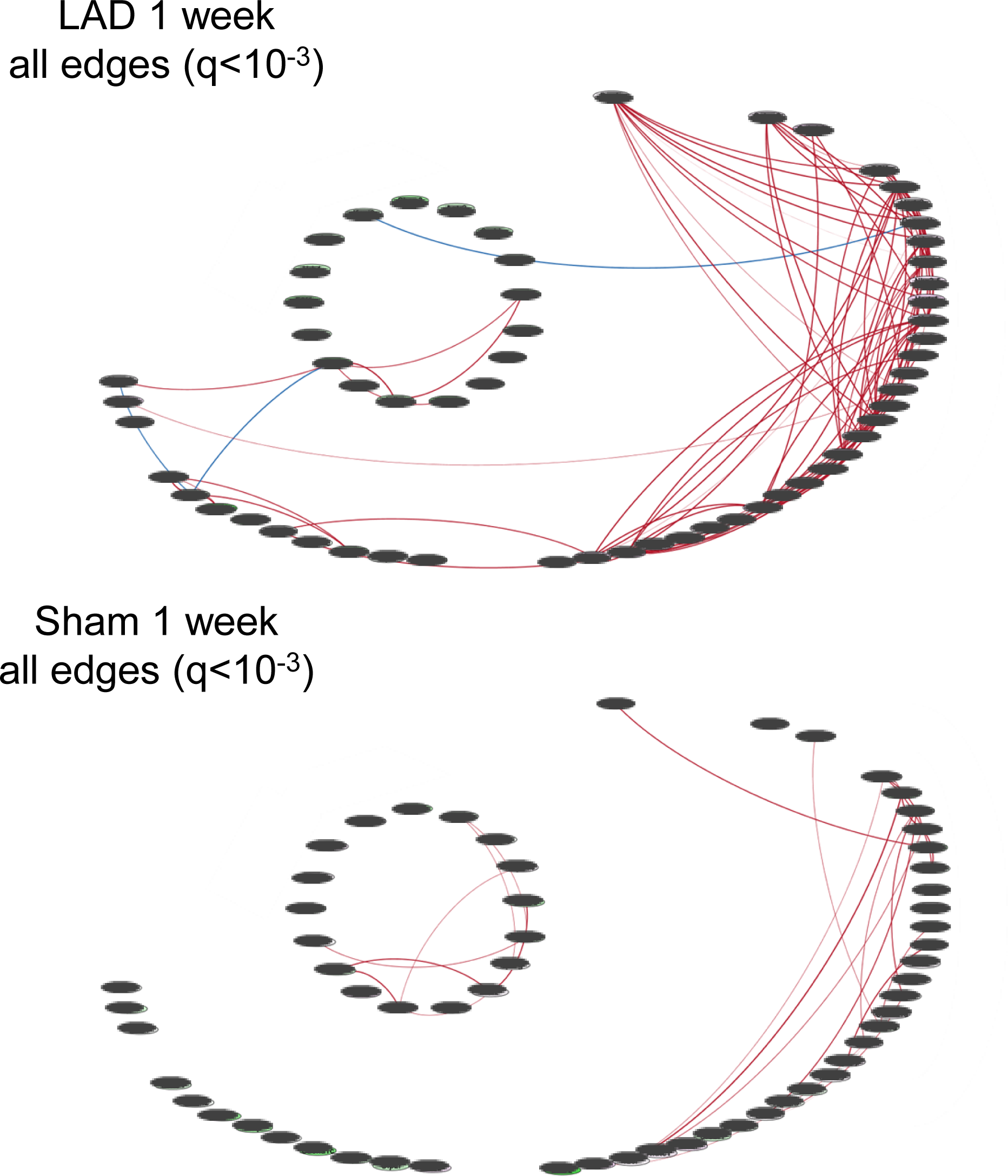
Correlation networks of each experimental group showing all edges. Edges are derived from Pearson correlations with cutoff of q<10^-3^. Blue edges are negative correlations and red edges positive with the edge thickness proportional to the correlation coefficient.

**Supplemental Fig. S19:**
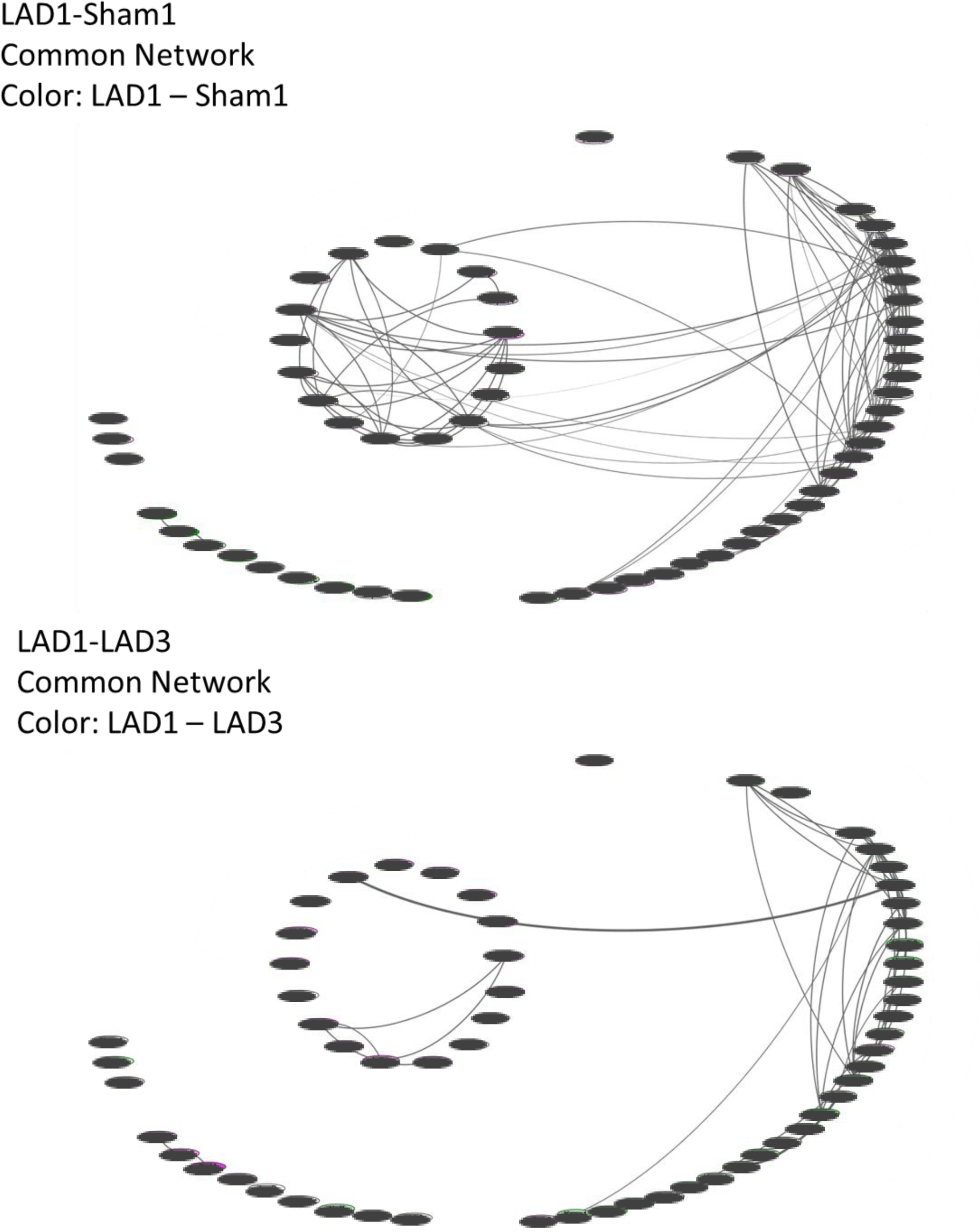
Network representations of common edges between two groups. Edge thickness is proportional to the average correlation coefficient. Inclusion of edges are from pearson correlations with q>0.001.

**Supplemental Fig. S20:**
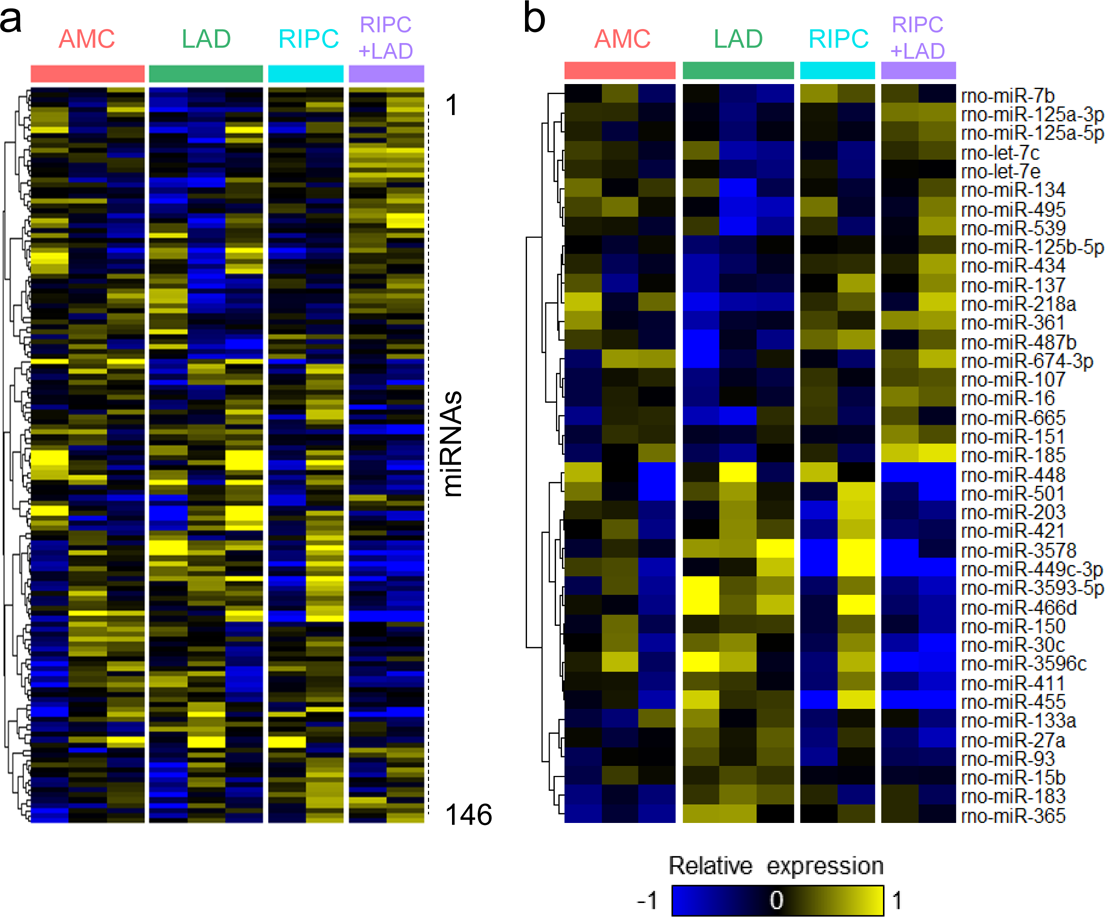
Expression profiles derived from Nanostring assays of microRNAs in whole DMV punches in response to LAD ligation with and without RIPC (see Fig. 4b). Samples are shown along the horizontal axis of the heatmap and microRNAs on the vertical axis.

**Supplemental Fig. S21:**
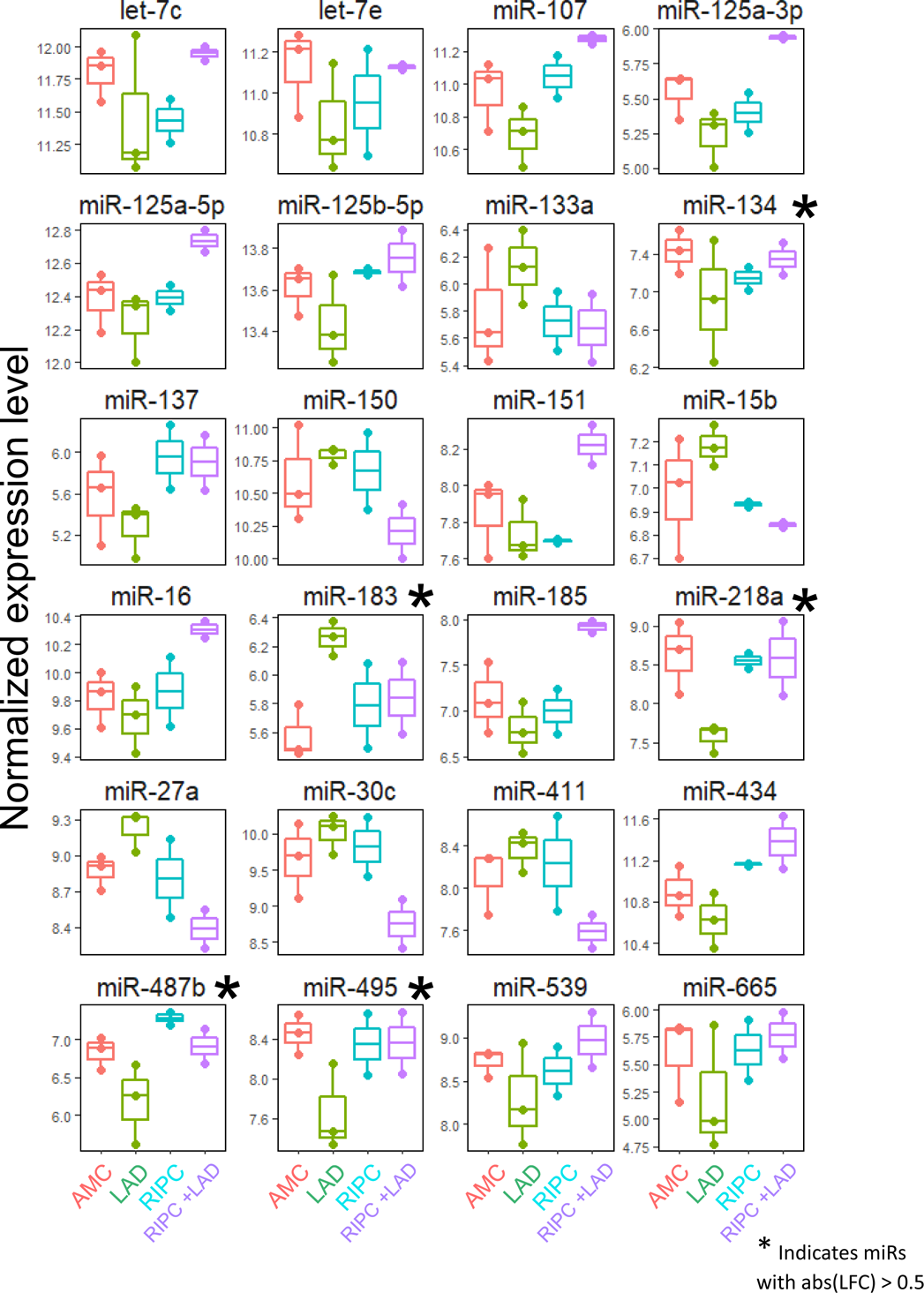
Expression profiles derived from Nanostring assays of microRNAs in whole DMV punches in response to LAD ligation with and without RIPC (see Fig. 4b). Box plots provide distribution of expression values for each experimental group.

**Supplementary Table S1:**
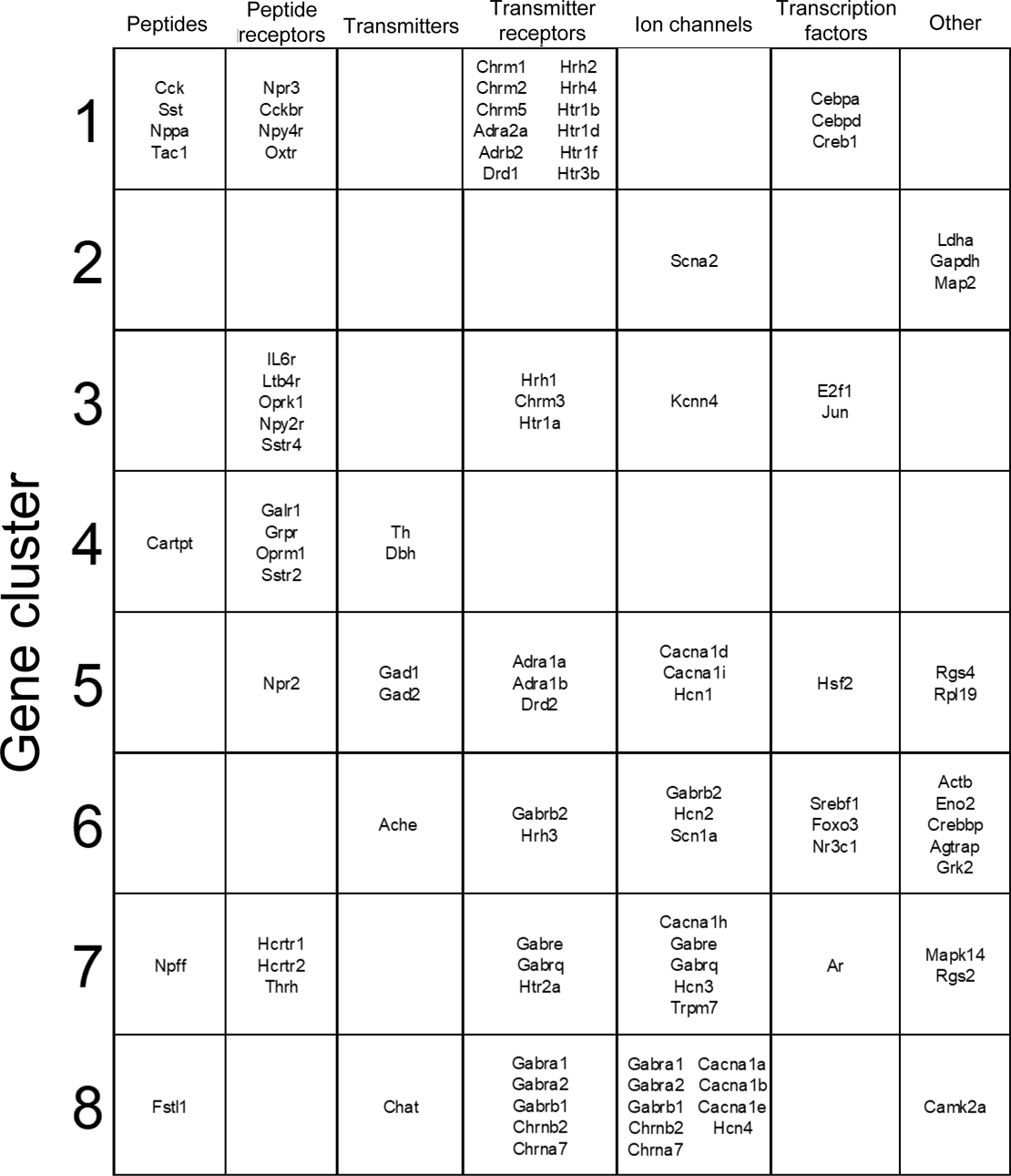
Categorization of genes from each of the clusters defined in the heatmap of Fig. 1B. Gene clusters were determined by unbiased hierarchical clustering, as described in methods. There are tendencies for ion channels to be in only a few clusters, suggesting that the milieu of channels expressed is coordinated rather than emergent from a series of diverse inputs.

**Supplementary Table S2:** Experimental design for chronic ischemia including distribution of sample collection and list of genes with categories.

| Surgery | Survival Time | Number of Animals | Number of Neurons | Number of Samples |
| --- | --- | --- | --- | --- |
| **Sham** | **1 week** | 3 | 133 | 19 |
| **LAD** | **1 week** | 3 | 136 | 17 |
| **Sham** | **3 weeks** | 3 | 158 | 21 |
| **LAD** | **3 weeks** | 4 | 279 | 34 |

| **<u>Ion Channels</u>** |  | **<u>Neuropeptides</u>** | **<u>Transcription Factors</u>** |  |
| --- | --- | --- | --- | --- |
| Cacna1a | Gabre | Agt | Agtr2 | Hsf2 |
| Cacna1b | Gabrq | Avp | Cebpa | Jun |
| Cacna1c | Grin2a | Cartpt | Cebpb | Jund |
| Cacna1d | Hcn2 | Cck | Cebpd | Maz |
| Cacna1e | Kcna1 | Gal | Creb1 | Mef2 |
| Gabra1 | Kcnj3 | Npff | Cutl1 | Nfatc1 |
| Gabra2 | Kcnn4 | Sst | E2f1 | Nfkb |
| Gabrb1 | Trpc6 | Tac1 | Fos | Nkx2-5 |
| Gabrb2 |  |  | Foxo3 | Pax4a |
|  |  |  | Foxo4 | Pparg |
|  |  |  | Hic1 | Srebf1 |
| **<u>G-protein Coupled Receptors</u>** |  |  | **<u>Modulators</u>** | **<u>Others</u>** |
| Adra1b | Drd1 | Hrh4 | Agtrap | Chat |
| Adra2 | Drd2 | IL6r | Camk2 | Ldha |
| Adrb2 | Drd4 | Ltb4r | Crebbp |  |
| Agtr1a | Fosb | Oxtr | Fstl1 |  |
| Chrm1 | Glp2r | Sstr2 | Grk2 |  |
| Chrm2 | Hcrtr1 | Sstr3 | Grk3 |  |
| Chrm3 | Hcrtr2 | Sstr5 | Mapk14 |  |
| Chrm4 | Hrh1 | Tnfrsf1a | Rgs3 |  |
| Chrm5 | Hrh2 |  | Rgs4 |  |

