## Supplemental Text for "Input-output signal processing plasticity of vagal motorneurons in response to cardiac ischemic injury"

**Summary of literature supporting the proposed input-output signal processing map of the DMV**

A great deal of effort has been expended to determine the inputs the DMV receives, both from an anatomical as well as molecular perspective. Some of these projections have also been characterized as to their effects on the physiology of DMV targets. In spite of this, connectivity information has not yet been collected into a processing map of the DMV as we have here in Figure 2. While this work has been important in deciphering the role of the DMV in coordinating and mediating CNS signaling. The following is a brief review of literature highlighting the axonal projections to the DMV and the neurotransmitters that are unique to each, therefore generating an input/output map of the DMV upon which the current data can be used to infer potential bandwidth of signaling from each source through receptor expression and effector transmitters and neuropeptides through enzyme and propeptide expression. There is the possibility that this map will require updating as more data becomes available, but this represents the state of knowledge at this current time.

*Afferent Projections to DMV*

The paraventricular nucleus (PVN) innervates the DMV with excitatory projections that contain combinations of several peptides, including oxytocin [1,2], bombesins including gastrin releasing peptide [3], atrial natriuretic peptide [4], cholecystokinin [5], and some sparse dopaminergic fibers as well [6]. These projections have been shown to mediate stress, satiety, and cardiovascular behavior. The lateral hypothalamus (LHA) sends projections containing orexins and brain natriuretic peptides [4,7]. Also from the hypothalamus are histaminergic tuberomammillary projections and synapse prominently in the DMV as well as the NTS [8–11]. The raphe pallidus (RPa) also receives projections from the PVN among other places, and sends efferent fibers to the DMV. These efferents are known to express serotonin and thyrotropin releasing hormone [12–14]. These projections have been shown to mediate both cardiovascular and gastrointestinal functions. The central amygdala (CeA) and bed nucleus of the stria terminalis (BST) send projections to the DMV containing corticotropin releasing hormone, neurotensin, tachykinins like substance P [15], and somatostatin [16]. There are also a large number of projections from the nucleus of the solitary tract and area postrema that include combinations of glucagon-like peptide 1, GABA, glutamate, norepinephrine, and dopamine [6,17]. There are excitatory adrenergic projections from the caudal ventrolateral medulla C1 population that terminate in the DMV [18] and the locus coeruleus [19]. Afferent cranial nerve nuclei send projections to the DMV including the trigeminal nucleus [20] and vestibular nucleus [21], both of which mediate physiological reflex activity of the cardiorespiratory system.

*Efferent Projections from DMV*

Along with canonical vagal efferent projections, there are some efferents that stay within the CNS, projecting to the parabrachial nucleus, cerebellar cortex, and deep cerebellar nuclei [22]. There is also a population of interneurons, described as GABAergic, that appear to project locally within the DMV as well as into the NTS [23]. Still the vast majority of DMV projections that leave the dorsal vagal complex run within the vagus nerve out to the periphery, innervating most major organ systems. Most prominent are gastrointestinal projections encompassing between 80-90% of all DMV-derived vagal efferent axons [23–25]. It should be noted that there are still a substantial number of axons that project to the intracardiac ganglia as well, about 5% of all DMV vagal efferents [26,27], roughly equivalent to the number of axons that originate from the NA [28]. Since the molecular understanding of such projections has been derived largely from pharmacological studies, a strong characterization of the peptidergic signaling targets has not been well-defined. It is known that DMV neurons express several neuropeptides, along with acetylcholine, dopamine, and nitric oxide. However, the target specificity of such projections still remains to be fully characterized. Some work toward this has described specific dopaminergic DMV projections to the pancreas [29], catecholaminergic projections to the gastric fundus [30], and nitric oxide, vasoactive intestinal peptide, and nicotinic receptors on gut macrophages that are signaled via the vagus nerve [31].

**Summary of literature for interpreting the neuromodulatory functions of DMV neurons**

*Gad+ DMV Neurons*

Almost all Gad+ single neurons express the canonical glutamatergic gene Camk2a [32] at some level. There is ample evidence for the existence of neurons that are simultaneously GABAergic and glutamatergic, even releasing both from the same terminals at the same time [33–39]. While this has been described at multiple regions of the rat brain, this is the first evidence we are aware of for the presence of such neurons in the dorsal motor nucleus of the vagus. It is surprising that nearly all neurons that express one or both of the glutamate decarboxylase genes co-express Camk2a. Almost all of the single cells assayed here expression Camk2a, with just less than half expressing Gad1 or Gad2. The implications for this are not entirely clear, but it suggests that is likely true that GABAergic neurons do project into the periphery, given that almost half of all single cells assayed at random can produce GABA. Given the plasticity that our work from the DMV in heart failure suggests, it may also be the case that the ability to generate GABA may come and go based upon the milieu of inputs driving these neurons. Further, GABA has been shown to stimulate release of GLP-1 [40], cholecystokinin [41], 5-HT [42], and gastric acid [43] in the gastrointestinal system. In Gad+ neurons, there are several that co-express Chat as well. There is a negative correlation between Chat and both Gad1 and Gad2 across all Gad+ cells. This suggests a gradient in the expression of these genes that may indicate a gradient in the ability to co-release GABA and acetylcholine. The presence of such a gradient lends credence to the ability of these neurons to switch from one phenotype to another.

This analysis suggests that there are distinct transcriptional states of GABAergic neurons in the DMV (Supplemental Figure S5). While four clusters are identified here, it is likely that some of these clusters may represent neurons of a similar state, but with different transient behaviors that might be further defined by dynamic studies. For purposes of this initial discussion, it is assumed that the four clusters represent four distinct functional states. There are several genes that show enriched or suppressed expression unique to each cluster and these may give clues to what makes each phenotype unique (Supplemental Figures XX-YY). For G_A the calcium channels Cacna1a and Cacna1i are suppressed along with Rgs2. Nppa expression is restricted almost entirely to G_A and Chrm5 and Kcnn4 are heavily enriched in G_A. For G_B, the most relatively suppressed genes include Gad1, Htr1a, and Kcna1. More notable are the enriched genes Chat and Fstl1 that are expressed in almost all neurons sampled from G_B and almost unexpressed in G_C. Also of note for G_C is the suppression of Adra1b. G_C is enriched for

Gad1 and Scn1a. While Cacna1i is suppressed in G_A, it is very strongly expressed in G_C and G_D. Of all the other genes uniquely expressed in G_D, most have no representation in G_C. This includes Adra1b, Fos, and Npff. Of note as well are Cacna1g and Dbh, both of which are starkly increased in G_D.

It may be possible to infer functional distinctions from each of the Gad+ clusters by contrasting their gene expression patterns. The most prominent feature of G_A is its unique expression of Nppa, which codes for the atrial natriuretic peptide (ANP), a protein heavily enriched in cardiac myocytes that is released to signal increased pressure within the heart [44] (Supplemental Figure XX). One of the other clusters, G_D, has enriched expression of the ANP receptor Npr2. Npr2 is also expressed in cardiac neurons as well as enteric neurons. Also of interest is the high expression of Kcnn4, the mediator of the $I_K$ potassium current that generally acts to maintain a hyperpolarized state. It may be that the hyperpolarized tendency is meant to increase the barrier to fire an action potential, requiring a strong depolarization to fire. Taken together, the GABAergic neurons of G_A are hypothesized to project out the periphery. They lack several calcium channels characteristic a burst or tonic firing phenotype and the expression of Nppa makes these neurons enticing as effectors in the periphery as does their enhanced potential for responsiveness to peptides like somatostatin and cholecystokinin based upon relative receptor expression.

Characterization of G_B includes unique downregulation of Htr1a in favor of Htr2a, representing a shift from an inhibitory response ($G_i$) to serotonin to an excitatory response ($G_q$). This comes along with a uniquely high expression of Chat and Fstl1 (SupplementalFigure XX). In some parts of the brain, cholinergic and GABAergic neurons are distinct populations [44], yet they have been shown to also potentially be one in the same [45]. Fstl1 codes for the protein follistatin-1, which apart from its well-described role during embryological development has been shown to exhibit robust cardioprotective effects against hypertrophy and ischemia-reperfusion injury, even so far as to induce myocyte regeneration in mammalian heart tissue [46,47]. Follistatin-1 is normally secreted from cardiac myocytes, but in the face of ischemia and myocyte death, this secretion is significantly reduced by the dying myocytes with others increasing their secretion [48]. Some evidence exists for replacement of follistatin-1 as an exogenous cardioprotective agent [46]. While this may make expression of Fstl1 a likely candidate for one of the mediators of the cardioprotective effects of the DMV, there is no significant upregulation observed in the IPC group. Although other work done in the DMV on a longer time course for heart failure does indeed demonstrate a substantial upregulation primarily three weeks after infarct and much less so at one week post-infarct. The high expression of Chat and Fstl1 make the GABAergic neurons of G_B another attractive candidate for projecting motor neurons from the DMV to the periphery. The other two subtypes defined here have qualities that are suggestive of a role as integrative interneurons rather than projecting effectors.

Perhaps more like canonical GABAergic neurons, G_C neurons (Supplemental Figure XX) have very low expression of Chat coupled with high expression of Gad1 and Scn1a, the latter of which has been shown to be essential for action potential firing more specifically in GABAergic neurons although it is present in several neuronal subtypes [49]. The high expression of Cacna1i is suggestive of a neuron that is capable of rebound burst firing at hyperpolarized resting membrane potentials [50–53]. This may suggest G_C neurons to be the most likely candidate of the four to be tonic firing interneurons although a great deal more work will be needed to verify this supposition.

Much like G_C, neurons in G_D also express higher levels of Cacna1i along with uniquely expressing high levels of Cacna1g, another low voltage gated T-type calcium channel implicated in burst firing at low resting membrane potentials [53]. Unique to this cluster is a larger percentage of neurons expressing higher levels of Adra1b, Dbh, Npff (Supplemental Figure XX). The $\alpha_{1B}$ receptor subunit encoded by Adra1b has been associated with susceptibility to synucleinopathy both in over-expression models as well as in *in vivo* studies [54,55]. While the implications of this are not clear at present it bears mentioning given the viable, yet debated, Braak hypothesis that places the DMV at the forefront of Parkinson's disease pathology preceding pathology in the substantia nigra [56–58]. All subtypes of GABAergic neurons examined here express some collection of adrenergic receptors, suggesting their ability to be influenced by projections from A2 neurons of the NTS or C1 neurons of CVLM among others. However, the more unique expression Adra1b in lieu of Adra1a, which is more highly expressed in G_B and G_C, may suggest a different response to adrenergic stimulation. The $\alpha_{1A}$ favors mediation of calcium influx whereas $\alpha_{1B}$ favors generation of inositol triphosphate as a second messenger [59]. As these neurons are effectively the only GABAergic neurons that also produce NE (due to Dbh expression) and Npff, they may be considered to be more sympathetic-like rather than parasympathetic-like.

There are two potential implications for neurons reliably expressing low-voltage activated T-type calcium channels (especially G_C and G_D that will be considered here. One is that they phenotypically maintain a low resting membrane potential so that they might flux calcium without firing action potentials. Such "subthreshold" calcium fluxes have been associated with a secretory phenotype where the oscillations are associated preferentially with peptide release rather than neurotransmitters [60–63]. However, it is also possible that the burst firing serves to aid in network synchronization as is the case in thalamic relay networks [53]. Through maintaining large windows of depolarization with windows of hyperpolarization, the "decision" to fire an action potential requires synchrony of inputs. This latter role for burst firing provides an excellent control concept for nucleus that has many neurosecretory cells with pacemaker activity [60,64]. This may provide one source by which vagal withdrawal is mediated, especially if the putative inputs from A2 and C1 cell populations are considered. These are neurons that are known to mediate sympathetic effects in their downstream targets in the periphery. If the sympathetic effectors fire with certain patterns, it is possible that they can activate GABAergic interneurons that might modulate DMV secretomotor neurons. This modulation does not necessarily mean inhibition, as a lowering of the resting membrane potential of neurons can both inhibit action potential firing as well as encourage subthreshold calcium flux, the latter of which may mediate peptide release in a secretory fashion.

The presence of GABAergic cell bodies in the DMV was once up for debate, but with improving techniques for protein detection and gene expression sensitivity, their presence has been confirmed [65,66]. There still exists a debate whether these GABAergic neurons act as local inhibitory interneurons, project out to the periphery, or both. With the characterization begun with this study it may be possible to define markers of projecting GABAergic neurons versus GABAergic interneurons, especially in light of the evidence mentioned previously for neurons that are a hybrid of canonical excitatory and inhibitory phenotypes. This concept and the evidence presented here with the blurred lines of distinction between GABAergic neuron subtypes strongly suggest a transcriptional plasticity of "terminally" differentiated neurons that is only beginning to gain broad acceptance.

*Fos+ DMV Neurons*

One of the underlying hypotheses of this study had been that the cardioprotective effects are being driven by changes in the activity of certain neurons in the DMV. Fos has been shown to be a reliable metric of recent neuronal activity [67] and serves to identify neurons that were active within the RIPC effect timeframe.

Upon further examination, it may be the case that Fos that does not accurately account for subthreshold ion flux activity nor for activity that does not result in changes mediating long-term-potentiation [68]. At least in the neurons sampled, there were no appreciable differences in the RIPC cohort as compared with the sham (Supplemental Figure S9). This may indicate that the effect is mediated by activity in such a small number of neurons that the sampling performed here was inadequate to detect them. More likely, however, this is due to the cardioprotective effects being mediated by changes in the molecular identity, like peptide expression, of the neurons that did not drastically alter activity. Another possibility is that the "activity" is not composed of strong depolarizations, but rather subthreshold fluxes. Regardless of the cause, this broadened the search to include all neurons sampled rather than just the Fos+ ones.

**Effects of sham surgery on shifting DMV neuronal states**

A surprising finding in this work was the prominent effect that the sham surgery had on gene expression in neurons of the DMV, even after just two and half hours. This is exemplified in the distribution of samples in the clusters shown in the heatmap in Figure 3. Several clusters differed from baseline in their sample distribution, most prominently State-s A2, C and D. Both A2 and C have an over-representation of sham samples whereas D has an over-representation of RIPC samples. The main differences from A1 to A2 can be found in parts of gene clusters 1,2, and 7 (Supplemental Table 1). The differences between the otherwise similar C and D clusters can be found mostly in gene cluster 1, the implications for which will also be discussed more specifically. In brief, they derive from RIPC neurons having expression patterns more similar to naive samples rather than sham, suggesting a repression of the stress response seen in sham. Given the projections to the DMV from several brain regions that are responsive to pain or stress, including PVN and CeA. The ability for certain neurons to change gene expression patterns so rapidly is indicative of their plasticity even as they respond to a brief surgical procedure under anesthesia.] When considering the effect of remote ischemic preconditioning, one of the noticeable features is an absence of the injury effect seen in sham, but not in the RIPC or the naive samples. There are numerous ways to generate a cardioprotective response, including pain and surgical stress [69]. However, others have reported a far more significant effect from peripheral ischemia as compared to a sham surgery as we performed in this study [70].

There is a large effect from the surgical treatment, observed in both RIPC and sham cohorts and not naive or observed uniquely in the sham cohort in the Fos+ samples (Figure S9). The most prominent is the significant upregulation of Th in samples from the RIPC and sham cohorts (ANOVA, p<0.001). It is only for Fos+ samples, the patterns of Th expression differ when all samples are taken into account as will be discussed.

The Sham response genes uncovered here include Cacna1g, Cckar, and Agtr1a. The definition of differential expression in this study includes a combination of either binomial testing (p<0.001) in assaying the percent of samples above the limit of detection and ANOVA (p<0.05 with Tukey post-hoc correction) for expression within the samples above the limit of detection (Figure 4A).

It is unclear what caused the increase in the low-voltage gated calcium channel Cacna1g (Cav3.1) in this context. It is known that this low conductance channel is involved primarily in pacemaking, calcium oscillations, and burst firing as are the other T-type channels [71]. Since the other T-type channels did not change in response either sham or RIPC treatment, it suggests that the firing patterns of these neurons will be different from the unchanged counterparts, but the nature of this change must be left to electrophysiological interrogation. The upregulation of Agtr1a and Cckar are associated with generalized stress responses and sympathetic drive in other contexts although their role in RIPC have not been described [5,72–74]. Such signaling in the DMV likely has its source in the NTS and CVLM with CCK signaling also potentially originating from neurons in PVN [75]. Indeed there is evidence for an upregulation of CCK production in PVN neurons in response to stress [5]. Using template matching, the genes that are most closely expressed in a similar pattern with Cckar include Th, Galr1 (galanin receptor 1), and Sstr2 (somatostatin receptor 2). Galanin signaling may also come from the NTS, but may also include signaling from the central amygdala (CeA) and bed nucleus of the stria terminalis (BST) [75]. Somatostatin projections have their origins most likely in the CeA and BST [16,76], a region well known to be one of the coordinators of stress in the CNS (de Kloet 2002). Galr1 has been shown to mediate stress responses in Th expressing neurons of other brain regions, like the locus coeruleus (LC) and may have similar effects in the DMV [77]. Delivery of galanin directly to the DMV has the net effect of causing hyperpolarization in gut-related DMV neurons [78]. The upregulation of Hrh3 also argues for decreased excitability owing to its ability to reduce the release of Ach and other neurotransmitters through presynaptic activity albeit without an appreciable difference in excitability [8,11]. Taken together, there is an overall sense of inhibiting normal gut function in the surgical response neurons as inferred from these changes, but that does not mean that the neurons modulating molecular changes in the gut, like secretory function, are also inhibited.

Interestingly, the neurons that have Agtr1a upregulated tend to coexpress Cacna1g. While the pattern of Galr1 has some overlap with these samples, Th, Sstr2, and Cckar do not. This suggests that there may be two distinct neuronal subtypes driving the surgical response. Instead, Cacna1g and Agtr1a have patterns of expression more similar to Hrh3, Hcn2, and Npff. Hrh3 is one of the genes that is diminished in RIPC back down to naive levels. Npff has the ability to generate a positive feedback loop inducing its own expression and activating neurons in an autocrine and paracrine fashion in the autonomic control regions of the brainstem [79] [80]. Npff mediates a sympathetic-like increase in blood pressure, heart rate, and contractility when centrally or peripherally administered [81,82]. It is also implicated in modulating morphine and ketamine analgesia [83,84]. Taken together, this cohort of neurons appear to be mediating a combination of anesthetic and stress-related responses in the DMV. That gene expression changes in the DMV less than three hours after 15 minutes of transient ischemia might be detected is a demonstration of how responsive the DMV is to peripheral afferent signals.

**Summary of literature for interpreting the differential expression of mircoRNAs during LAD ligation with and without RIPC**

miR-495 has been implicated in many inflammatory processes through its regulation of NOD2, where NOD2 can affect the expression of other pro-inflammatory cytokines and has also been shown to suppress NLRP3 inflammasome signaling, providing protection to cardiac microvasculature following ischemia/reperfusion injury[69]. Dysregulation of miR-495 has been implicated in processes involving motor neurons, such as ALS, and, interestingly, has been implicated in the downregulation of the GluA2 subunit of the Gria2 receptor [85]. This is in contrast to the stimulatory effects on Gria2 suggested to be associated with miR-218a. miR-218a, which has been found to be particularly enriched in neurites, stimulates the translation of the GluA2 subunit of the Gria2 AMPA glutamate receptor and has also been shown to target the RE-1 silencing transcription factor (REST), a transcriptional repressor of GluA2. Therefore, miR-218a enhances the strength of excitatory synapses by inhibiting the REST complex as well as enhances excitability through stimulation of GluA2 [86]. Taken together, the downregulation in LAD alone and subsequent recovery of both miR-218a and miR-495 with rIPC preceding LAD shows careful control of neuron excitability as well as inflammatory processes, and suggests that treatment with miR-mimics could provide some level of cardiac protection prior to LAD. miR-183 targets a wide range of genes that have both positive and negative effects on neuronal processes and inflammatory processes [87]. In particular, miR-183 has been found to modulate genes related to neuron hyperexcitability and inflammation including Nav1.3, Bdnf, and Trpv1, a voltage-gated sodium channel, a neuroprotective agent, and a non-selective ion channel [88]. This is consistent with upregulation of miR-183 in the DMV following LAD, as well as its downregulation as a protective effect provided by RIPC.

**Effects of persistent cardiac ischemia on gene co-expression network topology**

Gene co-expression networks were generated for each of the experimental groups using Pearson correlations and filtered using a q value cut-off of $q<10^{-3}$. The unique edges for each of the networks generated are given in Figures 7 with the connectivity in any of the experimental groups were excluded from the full network for each group given in Supplemental Figures S12, S13, and S14. Genes with no connectivity in any of the networks based upon the cut-off criteria mentioned previously were excluded from all the network figures. It is clear from these networks that the LAD-1 group has the greatest overall connectivity with over half of all edges being unique that group. Many of these unique edges are the negative correlation between genes in GC-1 and GC-5. Also of note are unique edges of interconnectivity within GC-1 and GC-5 respectively. The notable unique edges of the Sham 1 week group include several negative correlations with Foxo4, Camk2, and Drd4 in GC-3 with genes in GC-1. Also, there are several unique positive correlations to genes in GC-5 from other GCs as well as some unique interconnectivity within GC-5. The Sham-3 group is most notable for its very sparse connectivity.The LAD-3 group has much lower connectivity than either of the 1 week groups and also less unique edges. Most surprising is the emergence of Pax4a as a unique hub gene in its relationship to many GC-5 genes.

Within each of the treatment cohorts, there are several highly connected genes as is the baseline expectation for a biological network, suspected to have scale-free topology[89]. Overall, a few genes are highly connected across most of the cohorts, most notable being Cacna1b that is a member of the top five connected genes in each cohort. Cebpd is highly connected in the LAD-1 group and the LAD-3 group, but with many unique edges in the LAD-1 group. This suggests a role of Cepbd in mediating gene expression in the acute response to cardiovascular injury that persists to some extent beyond the acute phase.

Although the connectivity among the measured genes in the LAD-3 group is not as high as the either of the 1 week groups, there are still some inferences to be made from what is uniquely connected. There are three genes of interest here: IL6r, Ltb4r, and Pax4a. While IL-6 is a well-described pro-inflammatory cytokine, this is not to suppose that the effects of IL-6 binding its receptor in all contexts serves to increase inflammation. In neurons, the IL-6 receptor mediates anti-apoptotic signaling and can serve to protect against reactive oxygen species associated with metabolic stressors[90,91]. The levels of IL6r do not differ significantly between LAD-1, LAD-3 and Sham-3 (it is suppressed in Sham-1 relative to the others). However, high connectivity of IL6r in the LAD-3 group suggests that there is some signaling that is mediated by the protein product that is having an effect on expression of other genes, including Ltb4r and the pacemaker contributing channels Kcna1 and Hcn2. Similarly, the leukotriene B4 receptor is a canonical mediator of inflammatory processes, but has a different role in neurons of the central nervous system, promoting neurogenesis and neural differentiation from progenitor stem cells[92]. There is a great deal of evidence for a similar role for the IL-6 receptor in neurons of the CNS[93]. It may be possible that a cohort of genes associated with IL6r and Ltb4r expression are related to neural progenitor phenotype if stem cells were under consideration. However, these cells are neurons and were selected based upon strong NeuN protein expression as evidenced through immunofluorescence. Since NeuN is only expressed in neural progenitor cells after the neuron "fate" has been determined[94], it is highly unlikely that any stem cells of pre-neuronal phenotype were selected for this study if any do in fact exist in the DMV of adult rats. Therefore, it is possible that reactivation of some of the differentiating cell programming is used to regress the terminally differentiated neurons and permit them to undergo a phenotype shift. The third unique hub gene may give a clue to the nature of this phenotype shift, since Pax4a mediates differentiation into a neurosecretory phenotype both in the central nervous system as well as for the specialized secretory cells in the pancreas[95,96]. This is supported by upregulation of several genes in GC-5 that are coordinated by the three hub genes, IL6r, Ltb4r, and Pax4a. Many genes in GC-5 are suggestive of a neurosecretory phenotype, including several ion channels (Cacna1d, Kcna1, Hcn2)[60,64], neuropeptide signaling (Sstr2, Sstr5, Cck, Hcrtr1, Hcrtr2), and GABA receptors (Gabra1, Gabra2, Gabrb1, Gabrb2)[40]. While not verified directly in this work here, many of the genes that are well-connected n the unique LAD-3 network are those under the influence of the repressor element 1 silencing transcription factor (REST) complex, an essential regulator of a neurosecretory phenotype[97–99]. More work is needed to consider the possibility that repression of REST plays a role in the phenotype shift toward a neurosecretory phenotype.

**Effects of persistent cardiac ischemia on right versus left DMV neuronal gene expression**

A differential expression analysis was performed using a nested ANOVA to account for multiple measurements within each animal. While several genes did show differential expression between the sham and LAD groups at both 1 week and 3 weeks post-surgery, these significant differences can be best explained by changes on either the right side or left side only (Supplemental Figure S13). Of the six genes found to be significantly different, only Camk2 showed the bilateral difference of decreased expression in LAD-1 compared with Sham-1. This overall lower expression seems to be due to a loss of a bimodal distribution that is present in the sham condition. The other five genes from this study whose expression levels differed showed a pronounced effect on only the right or the left side (Supplemental Figure S13). Only Sst (somatostatin precursor) was shown to change only on the left side of the DMV with the largely increased expression reaching significance at the 3 week time point. The right side did show increased Sst expression at 3 weeks as well, but it did not rise to the level of statistical significance due to the more broad distribution of expression values. Four genes showed significant changes on the right side of the DMV: Hrh2, Npff, Fosb, and Cacna1e. The largest effect size here is the diminished expression of Npff in LAD-1 compared with Sham-1. Fosb showed decreased expression on the right only at the 1 week time point and Cacna1e showed increased expression only at the 3 week time point. Hrh2 expression had an interesting pattern, wherein the expression on the right in the sham condition at both time points was much higher than the left, which was at a comparable level to both sides in the LAD condition at both time points. The LAD ligation surgery was associated with a shift down to left sided levels at both the 1 week and 3 week points.
